# Somatostatin peptide signaling dampens cortical circuits and promotes exploratory behavior

**DOI:** 10.1101/2022.07.18.499817

**Authors:** Dakota F. Brockway, Keith R. Griffith, Chloe M. Aloimonos, Thomas T. Clarity, Brody Moyer J., Grace C. Smith, Nigel C. Dao, Md Shakhawat Hossain, Patrick J. Drew, Joshua A. Gordon, David A. Kupferschmidt, Nicole A. Crowley

## Abstract

Somatostatin (SST) neurons in the prelimbic (PL) cortex mediate a variety of behavioral states. However, little is known about the actions of somatostatin peptide signaling in shaping cortical functioning and behavior. Here, we sought to characterize the unique physiological and behavioral roles of the SST peptide in the PL cortex. We employed a combination of *ex vivo* pharmacologic and optogenetic electrophysiology, *in vivo* calcium monitoring, and *in vivo* peptide pharmacology to explore the role of SST neuron and peptide signaling in the mouse PL cortex. Whole-cell slice electrophysiology was conducted in pyramidal and GABAergic neurons in the PL cortex of C57BL/6J and SST-IRES-Cre male and female mice to characterize the pharmacological mechanism of SST signaling. Fiber photometry of GCaMP6f fluorescent calcium signals from SST neurons was conducted to characterize the activity profile of SST neurons during exploration of an elevated plus maze (EPM) and open field test (OFT). We further used local delivery of both a broad SST receptor (SSTR) agonist and antagonist into bilateral PL cortex to test causal effects of SST administration and receptor blockade on these same exploratory behaviors. SSTR activation broadly hyperpolarized layer 2/3 pyramidal neurons in the PL cortex in both male and female mice *ex vivo*, an effect that was recapitulated with optogenetic stimulation of SST neurons, through both monosynaptic and polysynaptic GABA neuron-mediated mechanisms of action. This hyperpolarization was blocked by pre-application of the SSTR antagonist cyclo-somatostatin (cyclo-SST) and was non-reversible. SST neurons in PL were activated during EPM and OFT exploration, indicating task-related recruitment of these neurons. Lastly, in line with this exploration-related activity profile, SSTR agonist administration directly into the PL enhanced open arm exploration in the EPM, while *in vivo* administration of an antagonist had no effect. Together, this work describes a broad ability for SST peptide signaling to modulate microcircuits within the prefrontal cortex and related exploratory behaviors.

## INTRODUCTION

The prefrontal cortex contains a complex microcircuitry of γ-Aminobutyric acid (GABA)-expressing inhibitory neurons capable of modulating excitatory cortical outputs involved in orchestrating a range of behaviors. Somatostatin (SST) neurons within the prelimbic (PL) cortex have been implicated in a variety of neuropsychiatric diseases and associated behavioral states^1, 2^. SST neurons play a role in binge alcohol drinking^3^, fear learning^4^, and the interaction between substance use and avoidance states^5^. These neurons also facilitate oscillatory synchrony between the prefrontal cortex and the hippocampus^6^, an established neural correlate of avoidance behaviors^7–10^. In addition, SST neurons are active during restraint stress^11^, highlighting the rapid activation of SST neurons during behaviors related to stress or threat. However, despite the abundance of this peptidergic neuronal subtype within the PL cortex^2^, little is known about how SST itself functions as a signaling molecule in this region, and what role this peptide plays in modulating behavior. Traditionally, it has proven difficult to separate GABAergic from peptidergic transmission due to technological barriers in probing these signaling molecules separately. Understanding the potential differing roles of GABA and SST is vital for understanding the unique contributions of SST to neuronal modulation, as well as for uncovering its potential as a therapeutic target.

Neuropeptides including SST are stored in dense core vesicles^12^, and can often diffuse greater distances than traditional neurotransmitters^13^. They exert long-lasting effects by activating G-protein coupled receptor (GPCR) signaling cascades. These properties of neuropeptides position them to modulate neuronal circuits and behaviors in unique and diverse ways^2, 14^ that can complement co-released GABA^2, 14^. SST signals through binding at five GPCRs that are predominately G_i/o_ coupled and are expressed throughout the mammalian cortex^15, 16^. The human clinical literature suggests a strong ‘pro-resiliency’ role for SST^2, 17^. SST mRNA is decreased in the post-mortem prefrontal cortex of individuals with bipolar disorder^18^, major depressive disorder^19^ and schizophrenia^20^. Similarly, recent clinical work demonstrated that alcohol-induced changes in local functional connectivity are dependent on overall SST gene expression in healthy individuals, with greater SST gene expression corresponding to decreased alcohol-induced changes^21^. Together, the human literature provides ample evidence for SST as a positive marker for the healthy brain, with decreasing expression, independent of GABAergic changes, associated with a host of neuropsychiatric disorders.

While we have previously characterized the activity conditions that promote endogenous release of SST from PL cortical neurons^22^, little work has systematically explored the downstream actions of SST. We demonstrate key actions of SST peptide signaling on modulating prelimbic cortical circuits, and promoting exploratory behaviors using a three-pronged approach. First, we used slice electrophysiology to demonstrate key physiological actions of SST peptidergic signaling on cortical microcircuits, using both exogenous bath application and stimulation of endogenous SST release with optogenetics. Second, we turned to *in vivo* fiber photometry to assess whether these neurons are activated during PFC-dependent exploratory behaviors in a manner that might promote endogenous peptide release. Lastly, we directly administered an SSTR agonist, and separately, an SSTR antagonist, into the PL cortex to assess the causal effects of SST signaling on these same exploratory behaviors.

Together, this work demonstrates that SST acts in the PL cortex to hyperpolarize cortical neurons, that SST neurons are recruited during exploratory behaviors, and that exogenous SSTR activation promotes exploration, suggesting that SST signaling is a promising target for modulating cortical circuits.

## MATERIALS AND METHODS

### Animals

All animal procedures were performed in accordance with the Institutional Animal Care and Use Committees (IACUC) at The Pennsylvania State University and The National Institute on Neurological Disorders and Stroke (NINDS), conforming to US National Institutes of Health guidelines. Adult male and female C57BL/6J mice were bred in house for electrophysiology or ordered from Jackson Labs (strain #000664, Bar Harbor, ME) for cannula experiments. For experiments involving *ex vivo* optogenetic activation of somatostatin cells, male and female SST-IRES-Cre^+/-^ on a C57BL/6J genetic background (#013044, Jackson Laboratory) were bred in house. Female and male VIP::Cre;SST::Flp mice (#010908 and 028579, respectively; Jackson Laboratory; heterozygous for both recombinases) were bred in house from crossings of homozygous VIP::Cre;SST::Flp females and Df(16)A^+/-^ males^23^ for the fiber photometry experiments. Df(16)A^+/-^ male breeders were backcrossed for more than 10 generations on a C57BL/6J background. All mice used for photometry were wildtype at the Df(16)A locus (i.e., Df(16)A^+/+^). Mice were maintained on a 12-h light cycle (lights on at 7:00 am, vivarium temperature 21°C, ±1°C) for all experiments. All electrophysiology and cannula experiments were conducted at The Pennsylvania State University, and all fiber photometry experiments were conducted at the NINDS.

### Electrophysiology

Mice were deeply anesthetized via inhaled isoflurane (5% in oxygen, v/v) and rapidly decapitated. Brains were quickly removed and processed according to the N-methyl-D-glucamine (NMDG) protective recovery method^24^. Brains were immediately placed in ice-cold oxygenated NMDG-HEPES artificial cerebrospinal fluid (aCSF) containing the following, in mM: 92 NMDG, 2.5 KCl, 1.25 NaH2PO4, 30 NaHCO3, 20 HEPES, 25 glucose, 2 thiourea, 5 Na-ascorbate, 3 Na-pyruvate, 0.5 CaCl2·2H2O, and 10 MgSO4·7H2O (pH to 7.3–7.4). The PL was identified according to the Allen Mouse Brain Atlas. 300-µm coronal slices containing the PL were prepared on a Compresstome Vibrating Microtome VF-300-0Z (Precisionary Instruments, Greenville, NC), and transferred to heated (31°C) NMDG-HEPES (in mM: 124 NaCl, 4.4 KCl, 2 CaCl2, 1.2 MgSO4, 1 NaH2PO4, 10.0 glucose, and 26.0 NaHCO3, pH 7.4, mOsm 300-310), for a maximum of 10 min. Slices were then transferred to heated (31°C) oxygenated normal aCSF where they were allowed to rest for at least 1 h before use. Finally, slices were moved to a submerged recording chamber where they were continuously perfused with the recording aCSF (2 mL per min flow rate, 31°C). Recording electrodes (3–6 MΩ) were pulled from thin-walled borosilicate glass capillaries with a Narishige PC-100 Puller. Drugs were included in the aCSF as described below per experiment.

Pyramidal and GABAergic neurons in layer 2/3 of the PL cortex were identified by location from midline, morphology (prominent triangular soma and apical dendrites for pyramidal neurons), and membrane characteristics, consistent with previously published electrophysiology in PL cortex layer 2/3 pyramidal neurons^3, 4, 25^. Pyramidal and non-pyramidal neurons were further confirmed by membrane properties and action potential width as appropriate^3^.

All experiments used a potassium-gluconate (KGluc)-based intracellular recording solution, containing the following (in mM): 135 K-Gluc, 5 NaCl, 2 MgCl2, 10 HEPES, 0.6 EGTA, 4 Na2ATP, and 0.4 Na2GTP (287– 290 mOsm, pH 7.35). Following rupture of the cell membrane, cells were held in current-clamp. A minimum of 5 min stable baseline was acquired prior to experiments and bath application of drugs. Measurements of intrinsic excitability were conducted at both resting membrane potential (RMP) and at the standard holding potential of –70 mV both before and after application of drugs. Gap-free RMP was recorded during the entire drug application period. Measurements of intrinsic excitability included the RMP, rheobase (the minimum amount of current needed to elicit an action potential during a current ramp protocol), action potential threshold (the membrane potential at which the first action potential fired during the current ramp), and the number of action potentials fired during a voltage-current plot protocol (VI plot). Rheobase and VI protocols were conducted consistent with previously published methods from our lab^26^. The rheobase protocol consisted of 4 sequential ramps, each injecting 120 pA of current, and each subsequent ramp stepping by 100 pA. Ramps lasted 1000 ms, and the protocol was stopped after the ramp during which the cell fired an action potential. The VI protocol consisted of increasing steps of depolarizing currents (0–200 pA, increasing by 10 pA per step, each step lasting 300 ms) with hyperpolarizing currents (not shown) included as a negative control. Some experiments were conducted in tetrodotoxin (500 nM) as noted to isolate monosynaptic activity. For all experiments where drugs were added to the aCSF, slices were perfused with drug as indicated per experiment, and slices were discarded after each experiment. Input resistance was monitored intermittently throughout each experiment, and when it deviated by more than 20% the experiment was discarded.

For optogenetic activation of SST cells with simultaneous electrophysiological recording of pyramidal cells in layer 2/3 of the PL cortex, slices were kept shielded from light, and experiments performed under low illumination. 25 µM Picrotoxin (Hello Bio, HB0506), and 1 µM CGP 55845 (Tocris, 1248) were added to the aCSF to block GABAergic signaling. A separate set of experiments were conducted with the addition of 1 µM cylcosomatostatin (cyclo-SST) in the aCSF along with GABA receptor antagonists. Cells were held in current-clamp and membrane potential (mV) was measured. Following establishment of a 5 min stable baseline, a 470-nm laser (CoolLED, United Kingdom) was directed to the slice for 10 min (10 Hz frequency). The membrane potential was recorded for an additional 5 min following stimulation. The perfusion pump was stopped before the start of the baseline file and remained off for the duration of the experiment to preserve released SST in the bath. SST cells in the PL were patched following the completion of a stimulation experiment and were stimulated at 10 Hz while recording membrane potential to ensure robust expression of ChR2. Following experiments, slices were immersed in 4% paraformaldehyde for 1 h, mounted on slides, coverslipped, and representative images of viral injection obtained on an epifluorescent microscope.

### Stereotaxic surgeries for cannula implantation

Custom bilateral cannulas targeted at the PL cortex were purchased from P1 Technologies (Roanoke, VA). Mice were deeply anesthetized with isoflurane (5% induction, 1-2% maintenance) and mounted on a stereotaxic frame (Stoelting, Wood Dale, IL). Following craniotomy, drill holes were targeted at the PL cortex (from Bregma: AP +1.8 mm, ML +/- 0.5 mm). Two additional drill holes were placed posterior to the injection site for bone screws. The guide cannula was lowered to the injection site for a final depth of -1.60 mm. Dental cement was used to secure the guide cannula, a dummy cannula was inserted, and Vetbond used for any additional scalp closure. Mice were allowed to recover, single housed, for a minimum of one week prior to behavioral testing. Placements of cannulas for all mice were verified by histology at the conclusion of the experiments.

### Stereotaxic surgeries for fiber photometry

Adeno-associated virus (AAV) encoding Flp-dependent GCaMP6f (AAV9.Ef1a.fDIO.GCaMP6f) was packaged by Vector Biolabs using a plasmid from Addgene (#118273). AAV encoding Cre-dependent TdTomato (AAV9.CAG.FLEX.TdTomato) was purchased from Addgene (#51503). Titers of the GCaMP6f and TdTomato viruses were 1×10^12^ and 1.9×10^13^ GC/mL, respectively, and were combined 10:1 (GCaMP:TdTomato) immediately prior to injection. All stereotaxic viral injections were conducted using aseptic surgical technique. Mice aged 12-16 weeks were deeply anesthetized with 5% isoflurane in oxygen (v/v) and secured in a stereotaxic frame (Kopf Instruments, Germany). Sedation was maintained using 1%–2% isoflurane during surgery. A midline incision was made on the scalp and two miniature screws (Antrin Miniature Specialities, Inc.) were secured to the skull. A craniotomy was performed above the left PL cortex according to the coordinates (in mm) +1.95 A/P (from bregma), -0.4 M/L (from bregma), -1.5 (from dura). The combined viruses were microinjected from pulled glass capillaries (using PC-100, Narishige) using a syringe (#7634-01, Hamilton) and syringe pump (#53311, Stoelting) with a volume of 500 nL and a rate of 100 nL/min. After infusion, the needle was left in place for 10 min to allow the virus to diffuse before the needle was slowly withdrawn. A fiber optic cannula (200µm core, 0.39 NA; CFMC12L05, ThorLabs) was implanted into the same craniotomy, with the coordinates (in mm) +1.95 A/P (from bregma), -0.4 M/L (from bregma), -1.0 (from dura). Dental cement (Unifast Trad) was used to adhere the ferrule/fiber to the skull. Tissue was secured to the dental cement with VetBond adhesive (1469Sb, 3M). Ketoprofen (5 mg/kg, SQ) was provided 30-min prior to the end of surgery for postoperative analgesia. Following surgery, mice were returned to group housing for 5-7 weeks prior to behavioral testing and photometric recording.

### Stereotaxic surgeries for optogenetic activation of SST cells

The viral construct AAV5-EF1a-DIO-hChR2(H134R)-eYFP-WPRE-HGHpA (Titer=2.1×10^13^ GC/mL) was obtained from Addgene (#20298). Male and female SST-IRES-Cre^+/-^ mice at least 8 weeks old were deeply anesthetized with 5% isoflurane and underwent stereotaxic surgery. Following craniotomy, mice were bilaterally injected using a 1 μL Neuros syringe (65458–01, Hamilton Company, Reno, NV; Stoelting) with 0.3 μL of the viral vector into the PL (AP: + 1.80, ML: ± 0.40, DV: − 2.30) at 0.1 μL/min. The syringe was left in place for 5 min to allow for diffusion before being slowly removed. Bupivicaine (0.1mL/20g) was applied topically and ketoprofen (0.1mL/10g) intraperitoneally for postoperative analgesia. Mice were allowed to recover for a minimum of 3 weeks prior to experiments to allow for adequate viral expression. PFC slices were visualized under infrared video microscope and blue LED (470 nm) for verification of viability and eYFP expression in the PL.

### In vivo fiber photometry apparatus

A custom-built spectrometer-based system (based on previously published systems^27–29)^ was used to conduct fiber photometry recordings. Blue light from a 473-nm laser (MBL-III-473-100mW, Ready Lasers) was directed through a Noise Eater (NEL01, ThorLabs) and into a kinematic fluorescence filter cube (DFM1, ThorLabs) onto a dual-edge dichroic mirror (ZT488/561rpc, Chroma). Light was then coupled using a FC/PC fiber coupler (PAF2-A4A, ThorLabs) into a fiber patchcord (M72L05, 200-um core, 0.39 NA, ThorLabs) connected to a fiberoptic rotary joint (FRJ_1×1_FC-FC, Doric Lenses) followed by another patchcord (200-um core, 0.39 NA, Doric). Blue light power was approximately 80 µW at the ferrule end of the final patchcord, resulting in ∼70 µW output from the surgically implanted ferrule. On each recording day, the surgically implanted ferrule was cleaned with 70% ethanol and lens paper (806, Ted Pella) and securely attached to the ferrule end of the final patchcord via a mating sleeve (SM-CS1140S, Precision Fiber Products). Fluorescence emission from the tissue was collected by the same fiber, filtered through a dual-band emission filter (ZET488/561m, Chroma), and directed using a fiber coupler (PAF2S-11A, Thorlabs) into a 200-µm core, anti-reflection-coated fiber (M200L02S-A, ThorLabs) which led to a spectrometer (Ocean Insight, QEPro). The spectrometer quantified photon counts across a ∼350-1130 nm wavelength window. Spectra comprised of integrated photons captured over a 37-ms time window were sampled at a frequency of 20 Hz.

### Drugs

For electrophysiology, Octreotide Acetate was dissolved in ddH2O at 3.27 mM, aliquoted at 100 µL, stored at -20°C, and diluted to 3.27 µM as needed. SST (Bachem, H-1490) was dissolved in ddH2O at 1 mM, aliquoted at 50 µL, stored at –20°C, and diluted to 1 µM in aCSF as needed. Cyclosomatostatin (cyclo-SST; Abcam, ab141211) was dissolved in DMSO at 1 mM, aliquoted at 100 µL, stored at –20°C, and diluted to 1 µM in aCSF as needed. Tetrodotoxin (TTX) (Abcam, ab120054) was dissolved in ddH2O at 5 mM, aliquoted at 50 µL, stored at –20°C, and diluted to 500 nM in aCSF as needed. 3 mM Kynurenic acid (Sigma, K3376), 25µM Picrotoxin (Hello Bio, HB0506), and 1µM CGP 55845 (Tocris, 1248) was added to the aCSF as needed. For behavior, Octreotide Acetate (Sigma Aldrich, PHR 1880) was dissolved in sterile aCSF at 327 µM, aliquoted at 100 µL, stored at –20°C, and diluted as needed.

### Drug microinjection procedure

Drug microinjection protocols were adapted from previously published studies^30^. Mice were habituated to handling and manipulation of the dummy cannula for 3 consecutive days prior to behavioral testing. Octreotide or aCSF control was injected at an infusion rate of 100 nL/min over 3 min. The infusion cannula was left in place for 2 min to allow for local diffusion of the injected solution. For behavior using administration of the SSTR-antagonist cyclo-SST, mice were habituated as described above. Similarly, cyclo-SST or aCSF control was injected at an infusion rate of 100 nL/min over 3 min and allowed to remain in place for 2 min to allow for local diffusion of the injected solution. Cyclo-SST was administered at a concentration of 0.01 µg / 300nL per hemisphere.

### Behavior

Identical open field test (OFT) and elevated plus maze (EPM) arenas were constructed at The Pennsylvania State University and NINDS. Behavioral experiments were conducted during the light cycle and mice were brought to the testing room and allowed to rest for at least 1 h prior to experimentation. Behavioral tests were separated by 48 hr and their order was counterbalanced across mice.

For the OFT, mice were initially placed in the corner of a 50 x 50 x 20 cm arena and allowed to explore for 30 min. One mouse jumped out of the OFT and was excluded from analysis. The total distance traveled over 30 min and the time spent in the 30 x 30 cm center square were quantified. The total distance traveled was displayed both as a time course (5-minute bins) and as a total value.

For the EPM test, mice were placed into the center square of an elevated (40 cm) crossbar with two open and two closed arms (30 x 5 cm), facing a closed arm (20 cm walls of Plexiglass). Mice were allowed to explore the maze for 5 min (drug cannula experiments) and 10 min (fiber photometry) and behavior was video recorded. Mice that fell off the EPM were excluded from the analysis (5 mice total). Percent time spent in the open arms, closed arms, center zone, and the percent of entries (open arm entries/total entries x 100) into the open arms were analyzed.

### Behavior with fiber photometry

Behavioral testing with photometry recordings was conducted as described above. Mice were first habituated to being handled and tethered to an optical fiber in a bucket for two daily 1-h periods prior to the first OFT/EPM test day. Each test day began with 10 min of blue light administration (∼70 µW) in the homecage to stabilize basal GCaMP6f fluorescence and further habituate each mouse to being tethered. A Python-controlled waveform generator (PulsePal v2, SanWorks) was used to simultaneously deliver 20-Hz TTLs to the spectrometer (to trigger spectral integration events) and camera (FLIR Blackfly S USB3; to trigger the camera shutter). Video frames generated at 20Hz were processed in Bonsai operating real-time DeepLabCut processing nodes^31, 32^ such that each frame was assigned coordinates for the maze and mouse as it was captured. Camera shutter events (also 20Hz) were simultaneously captured as digital events in OpenEphys to facilitate subsequent alignment of photometry and positional data (see Behavior Data Analysis).

Photometry/position data were recorded for 10 min during homecage behavior and throughout the subsequent EPM/OFT test. Recordings in both homecage and OFT/EPM were conducted in 2.5-min bins separated by 2 sec. These 2-sec gaps enabled brief openings of a Python-controlled clamp on the fiber optic rotary joint. The clamp prevented light artifacts due to rotary joint movement during recordings. These brief openings allowed for any accumulated patchcord tension to be released prior to reclamping and resumed recording.

### Histology for cannula/fiber optic placements and viral expression

Mice were deeply anesthetized with Avertin (250 mg/kg) or isoflurane (5%) and underwent transcardial perfusion, first with ice-cold phosphate buffered saline (PBS) followed by 4% (w/v) paraformaldehyde (PFA). Following perfusion, brains were post-fixed in PFA for 24 h, transferred to PBS and sectioned at 40 µm using a Leica vibratome (VS 1200, Leica) or at 50 µm using a Campden Instruments vibratome (7000 smz2). Sections were mounted on SuperFrost or Marienfeld UniMark glass slides, air dried, and then cover-slipped with Immu-Mount (Thermo Fisher Scientific, Waltham, MA, United States) or DAPI Fluoromount-G (Southern Biotech) mounting media. Slides were then imaged on an Olympus BX63 upright microscope (Center Valley, PA) or a Leica customized epifluorescence scope fitted with a CMS GmbH CMOS camera (Leica Microsystems). The image of representative fiber optic placement and viral expression was captured using a Zeiss LSM 800 confocal microscope. Areas containing the most damage were considered the central location of guide cannula or fiber optic placement. Damage location was then determined in reference to the Allen Mouse Brain Atlas and a 0.5-0.55 mm projection was added to the end of the damage to account for the internal cannula projection. Mice with cannula/fiber placement outside of the PL cortex were excluded from behavioral/photometry data analysis. 8 of 120 mice for cannula experiments (including mice used for pilot dose response curves, data not shown), and 1 of 12 mice for fiber photometry, were removed for misplaced implants.

### Behavior data analysis

Behavior for both cannula drug administration and fiber photometry were tracked with DeepLabCut^33^. Analysis of cannula drug administration assays was performed in MATLAB; analysis of photometry experiments was performed in Python. Experimenters were blinded to drug injection group throughout the behavioral testing and data analysis.

### Cannula microinfusion behavioral analysis

Behavior for cannula drug administration was tracked with DeepLabCut (DLC; Version 2.2.1) and analyzed with custom MATLAB algorithms. For OFT and EPM, three body positions were tracked for behavior classification: the head, middle of the body (caudal to the shoulder joint), and the trunk of the tail. Behavioral recordings that consisted of separate zones included additional tracked points bounding zones of interest. For OFT, these 16 points created boundaries that outlined the edge of the arena (1 point per corner of the 50 cm x 50 cm arena), the center zone (1 point per corner of the central 30 cm x 30 cm square), and each corner (1 point per corner of the 10 cm x 10 cm corner box in the outer zone). Recordings of EPM behavior included the tracking of 12 distinct points that outlined each arm of the maze. Any points that bounded the corner of two zones were only quantified with 1 distinct point to eliminate any redundancies in analysis. Recording of all cannula drug administration behavior was performed using a camera at a frame rate of 29.97 frames per second. OFT and EPM tracking utilized transfer learning to retrain residual network (resnet101) using 9 mice and 10 mice for each behavior, respectively. Mice used for training were selected from recordings performed on separate days whenever possible. DLC model training was considered adequate when confidence in the position was >95%, with most points achieving a confidence >99%. Tracked position coordinates were exported from DLC as CSV file for MATLAB analysis when appropriate confidence levels and tracking performance were reached.

OFT distance was classified as the Euclidean distance of the middle of the mouse body between subsequent frames. Time was quantified by the number of frames a mouse was in a polygon bounded by the corners of the zone of interest. All data were then converted from units of pixels/frames to cm/s using known bounds of the arena and the frame rate of the camera. Percent time in a specific zone was quantified as the total time in the zone divided by the total testing time.

EPM distance was classified as the Euclidean distance of the middle of the mouse body between subsequent frames. Time was initially quantified by the number of frames a mouse was in a polygon bounded by the corners of the zone of interest. All data were then converted from units of pixels/frames to cm/s using known bounds of the arena and the frame rate of the camera. Percent time in a specific zone was quantified as the total time in the zone divided by the total testing time. The central zone connecting arms was classified as a ‘dead zone’ and not included as a portion of the open or closed arms. Head dips were classified as an extension of the head beyond the bounds of the open arm. Random videos were selected, and behaviors were hand scored where possible to verify both tracking and algorithm accuracy. Experimenters were blinded to injection condition throughout the behavioral analysis.

### Photometry behavior analysis

Behavioral analysis for the photometry experiments was conducted using many of the same procedures and parameters as described above. In these experiments, Bonsai-DLC was used for behavioral tracking. Bonsai-DLC enabled use of a pre-trained DLC model in a Bonsai workflow to process live-streamed videoframes and generate DLC coordinates in real time. DLC models for the EPM and OFT were created by labelling 300-500 frames per test, comprised of approximately 10-20 frames from each of 20-30 videos of different mice with comparable surgeries/fiber optic tethers to the present experimental mice recorded on different days. In addition to the maze boundaries, six mouse body parts were labelled on each frame: nose, headcap, shoulder, midpoint, hind and base of the tail. The hind label was used for behavioral analysis. The model was trained for approximately 750,000-1,000,000 iterations, yielding confidence values of >99% in most cases.

A Python-controlled waveform generator (PulsePal v2, SanWorks) delivered 20-Hz TTLs to a FLIR Blackfly S USB3 camera. Each resulting frame was processed for all model-labelled body/maze parts. A confidence threshold of >95% was applied in Bonsai to the positional data. Exported data were down-sampled to 10Hz, and a Kalman Filter (pykalman.github.io) was applied to estimate position data for missing values with confidence <95%. The “opencv homography” Python function was applied to align cohorts with slight variations in camera angles. The hind position from each frame was assigned a maze zone based on the coordinates of the maze boundaries. Brief departures from an assigned zone (e.g., changes from “open_top” > “center” > “open_top”) of 3 or less frames were corrected (e.g., converted to all “open_top”) to account for noise in the detection around zone boundaries. In the OFT, the timing and number of zone transitions were then calculated based on these assigned zones. In the EPM, because transitions between arms were complicated by the intervening center zone, zone transitions were defined using two sets of criteria. In cases when changes in assigned zone occurred from one zone to another and back to the original (e.g., “open_top” > “center” > “open_top”), to be assigned a zone transition, the departure (e.g., to “center”) needed to last for a minimum of 6 frames. In cases when changes in assigned zone occurred from one zone to another to yet another (e.g., “open_top” > “center” > “closed_left”), to be assigned a zone transition, the departure (e.g., to “center”) needed to last for a minimum of 2 frames. These thresholds were implemented to prevent overcounting of re-entries into the same zone, and to accommodate counting of fast transitions between zones (e.g., rapid movement across the center from open to closed arms). Thresholds were determined by cross-referencing DLC-based assignment of transitions with manual assignment by an experimenter in several videos.

### Photometry data analysis

The shape and amplitude of the spectrometer-derived fluorescence spectra were used to confirm *in vivo* GCaMP6f and TdTomato expression. To separate the fluorescence derived from GCaMP6f and TdTomato, all raw emission spectra were transformed using a spectral linear unmixing algorithm written in R, as described previously ^27, 29^. The resulting unmixed GCaMP6f and TdTomato coefficients were linear regression-corrected to remove gradual reductions in signal due to fluorophore signal fading across the behavioral test. To control for movement artifacts in the fluorescence signal, the ratio of the unmixed GCaMP6f and TdTomato coefficients was calculated (GCaMP:TdTomato; G:T; as in^27^). The G:T timeseries was down-sampled from 20Hz to 10Hz. Z-scores of the G:T ratio were then calculated for each 10-Hz time point, using the mean and standard deviation of their corresponding 2.5-min bin of recorded data. Sample traces, their z-scored values, and the z-scored ratio are presented in **Supplemental Figure 5**. Once aligned to positional data from DeepLabCut, G:T z-scores were then averaged across task-relevant periods (e.g., open-vs. closed arms; center vs. surround zones), or aligned to discrete transitions (e.g., center-to-open arms; center-to-surround zones). G:T z-scores were averaged across all events (bouts in an arm/zone, or transitions) within a single animal, and then reported as the average z-score across animals.

Correlational analyses were performed to compare the degree to which photometry signals recorded during open vs. closed arm entries correlate with concurrent speed of the mice across these transitions. Using GraphPad prism, 1^st^ derivatives of the photometry and speed time series were calculated for all open and closed arm entries performed by all 11 mice. These derivative time series were averaged within mice and normalized to the peak value in a 4-sec peri-transition time window. Using the 40 0.1-sec timepoints in this 4-sec window (corresponding to the 10 Hz photometry and speed recordings), correlations were performed for each mouse, and the correlation coefficients were then averaged to assess whether the speed-photometry correlations were stronger during open or closed arm transitions.

### Statistics

Experimenters were blinded to group designations wherever possible until analysis was completed. Data were analyzed by one-sample t-tests, unpaired t-tests, paired t-tests, 2-way ANOVAs, 3-way ANOVAs, and post-hoc tests as appropriate and indicated for each experiment. Statistics and figure preparation were conducted in Prism 9 (GraphPad, La Jolla, CA). Data are expressed as means ± SEM for all figures and considered significant if *p* < 0.05. No statistical outliers were removed from the electrophysiology, photometry, or behavioral data.

## RESULTS

### Somatostatin has an inhibitory effect on PL cortical circuits

#### SST hyperpolarizes and decreases excitability of PL cortical pyramidal neurons

We first sought to understand what, if any, effect SST peptide signaling had on the main output neurons of the prefrontal cortex, glutamatergic pyramidal neurons. To examine the effect of SST on PL cortical circuits, *ex vivo* whole-cell current-clamp recordings were performed on pyramidal neurons in layer 2/3 of the PL cortex in adult male and female C57BL/6J mice (representative circuit diagram in **Figure 1A**). Layer 2/3 pyramidal neurons, a direct monosynaptic target of SST cells, were chosen to correspond with our previously published work on SST neurons in the PL cortex^3, 5, 22^. SST cells synapse onto pyramidal cells in layer 2/3 of the PL cortex^22^. Measurements of intrinsic excitability were conducted at both RMP and at the standard holding potential of -70 mV before and after 10 min bath application of 1 µM SST. 1 µM SST was chosen to correspond with previously published work on SST in cortical neurons^34^. Males and females were initially statistically analyzed separately. As no sex differences emerged in the overall effect of SST on pyramidal neurons, electrophysiology data are presented with the sexes combined. Representative traces of rheobase recordings are shown in **Figure 1B**, before and after SST application. SST hyperpolarized pyramidal neurons and significantly decreased the RMP (*t*_30_ = 3.679, *p* = 0.0009; **Figure 1C**). SST also significantly increased the rheobase (the minimum amount of current required to elicit an action potential) at both RMP (*t*_30_ = 6.571, *p* < 0.0001) and –70 mV (*t*_30_ = 4.105, *p* = 0.0003; **Figure 1D, 1F**). SST did not significantly alter the action potential threshold at RMP (*t*_30_ = 1.568, *p* = 0.1273) or –70 mV (*t*_30_ = 0.8883, *p* = 0.3814; **Figure 1E, 1G**). Representative traces of VI recordings are shown in **Figure 1H, 1I**. SST significantly reduced the number of action potentials fired in the VI plot at RMP (2-way ANOVA; *F*_current_(20,600) = 31.51, *p* < 0.0001; *F*_drug_(1,30) = 17.50, *p* = 0.0002, *F*_current x drug_(20,600) = 15.51, *p* < 0.0001; **Figure 1H;** significant post-hoc Bonferroni’s are indicated on figures). As SST affects the minimum amount of current needed to elicit an action potential (rheobase) but not the action potential threshold this suggests that SST is not acting on voltage gated sodium channels. There was also a significant current x drug interaction in the number of action potentials fired in the VI plot at –70 mV (2-way ANOVA; *F*_current_(20,580) = 24.22, p < 0.001; *F*_drug_(1,29) = 1.637, *p* = 0.2109, *F*_current x drug_(20,580) = 3.247, *p* < 0.0001; **Figure 1I;** significant post-hoc Bonferroni’s are indicated on figures). Several different mechanisms are likely occurring simultaneously (ex. GIRK channel activation^34^) as well as presynaptic changes in glutamate or GABA signaling which account for SST mediated hyperpolarization and decrease in intrinsic excitability (discussed in greater detail in the discussion).

**Figure 1.**
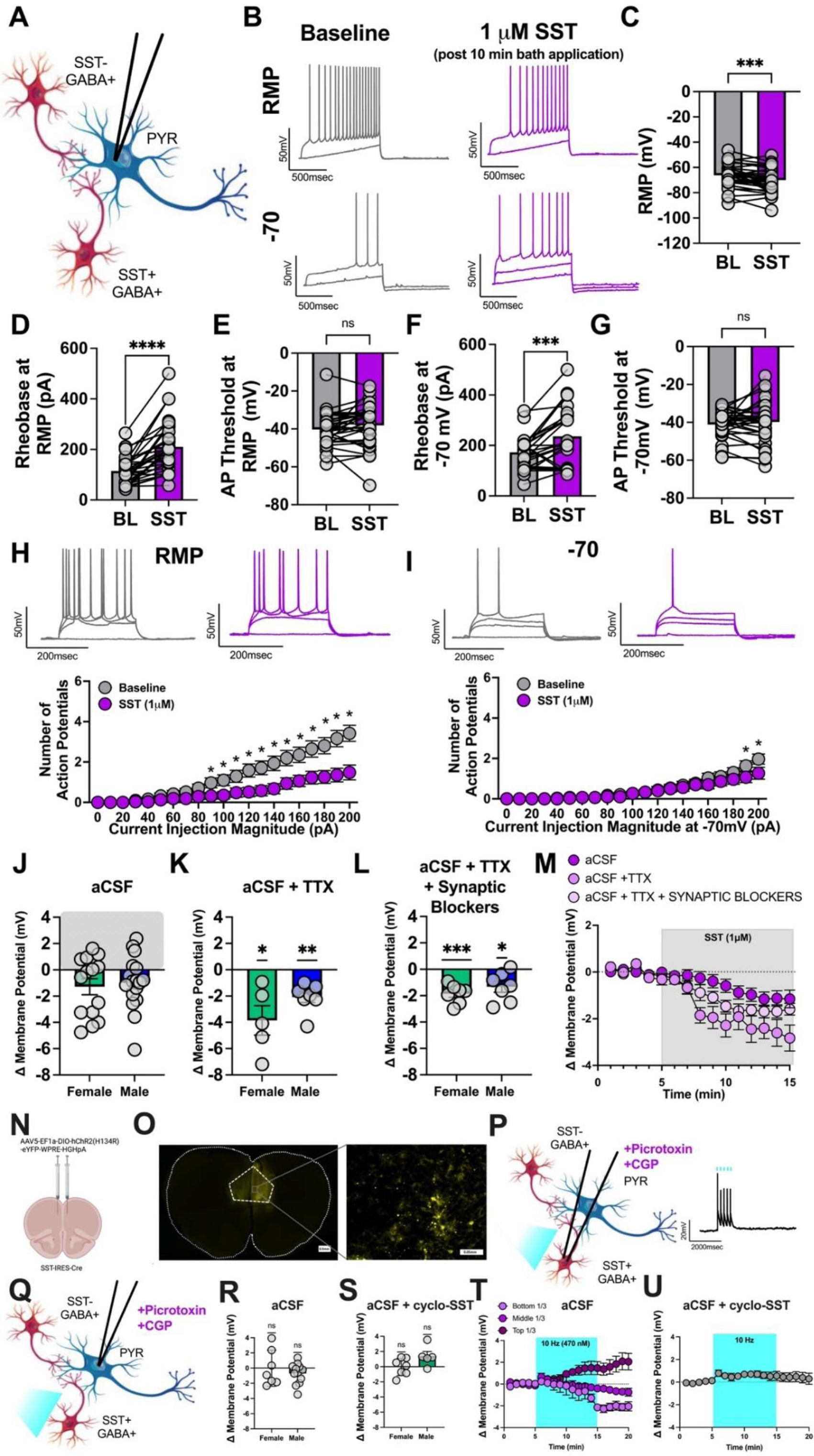
Somatostatin hyperpolarizes PL pyramidal neurons of male and female mice. **(A)** Schematic of experimental setup. Whole-cell current clamp recordings were conducted in PL cortex layer 2/3 pyramidal neurons in males and females. **(B)** Representative traces before (grey) and after (purple) 1 µM SST application at both RMP (top) and –70 mV (bottom) for rheobase experiments. **(C)** SST significantly decreased the RMP, and **(D)** significantly increased the rheobase at RMP. **(E)** SST did not significantly alter the action potential threshold at RMP. **(F)** Similar effects are seen at the common holding potential of –70 mV, with SST bath application significantly increasing the rheobase. **(G)** SST did not significantly alter the action potential threshold at -70. **(H)** Representative VI traces (corresponding to 0, 110, 150, and 200 pA of current) at RMP before (grey) and after (purple) SST application. SST significantly reduces the number of action potentials fired in response to increasing amounts of current injection at RMP. **(I)** Representative VI traces (corresponding to 0, 110, 150, and 200pA of current) at –70 mV before (grey) and after (purple) SST application. Significant reductions in action potential firing were seen at the highest current injection magnitudes, at the common holding potential of –70 mV. For panels A-H n=31 cells from 12 female and 13 male mice; for panels I n=30 cells from 11 female and 13 male mice. **(J)** Shift in membrane potential in aCSF (female in green, male in blue) following 10 min SST administration. For panel (J) n= 31 cells from 12 female and 13 male mice. **(K)** Shift in membrane potential (female in green, male in blue) following 10 min SST administration with the addition of 500nM TTX in the aCSF. For panel (K) n= 12 cells from 4 female and 5 male mice. **(L)** Shift in membrane potential (female in green, male in blue) following 10 min SST administration with the addition of 500nM TTX to block action potentials and 3 mM Kynurenic acid to block glutamate receptors, 25µM Picrotoxin to block GABA_A_ receptors, and 1µM CGP to block GABA_B_ receptors. For panel (L) n= 14 cells from 5 female and 6 male mice. **(M)** Data from panel J-L graphed as a function of time throughout the recording. **(N)** AAV5-EF1a-DIO-hChR2(H134R)-eYFP-WPRE-HGHpA was injected into the PL of adult SST-IRES-Cre +/-mice and was allowed to express for a minimum of 3 weeks. **(O)** Representative images of ChR2 injections. Left, ChR2 injections were confined to the prelimbic cortex (scalebar = 0.5mm 4x magnification). Right, 20x magnification of injection site and patching region (scalebar = 0.05mm). **(P)** SST neurons in the PL were patched and reliably elicited action potentials at 10Hz. 25µM Picrotoxin was added to the aCSF to block GABA_A_ receptors, and 1µM CGP to block GABA_B_ receptors. **(Q)** Pyramidal cells in the PL were patched and following a 5 min stable baseline SST cells were stimulated for 10 minutes (10Hz). 25µM Picrotoxin was added to the aCSF to block GABA_A_ receptors, and 1µM CGP to block GABA_B_ receptors. **(R)** Shift in membrane potential (female in green, male in blue) following 10 min SST activation (10 Hz) with the addition of 25µM Picrotoxin to the aCSF to block GABA_A_ receptors, and 1µM CGP to block GABA_B_ receptors. For panel Q, n= 19 cells from 4 female and 5 male mice. **(S)** Shift in membrane potential (female in green, male in blue) following 10 min SST activation (10 Hz) with the addition of 1µM cyclo-SST to block SST receptors, 25µM Picrotoxin to the aCSF to block GABA_A_ receptors, and 1µM CGP to block GABA_B_ receptors. For panel R, n= 12 cells from 5 female 3 male mice **(T)** Data from panel Q graphed as a function of time throughout the recording. Data was separated into thirds depending on the level of hyperpolarization following 10 minutes and graphed separately for visualization. Those that hyperpolarized the most are labeled bottom 1/3 and are depicted in light purple, middle 1/3 in medium purple and top 1/3 in deep purple. **(U)** Data from panel S graphed as a function of time throughout the recording.

Interestingly, meaningful variability in SST-induced changes in pyramidal neuron membrane potential emerged (**Figure 1J** for individual data and separated by sex). While SST hyperpolarized most pyramidal neurons recorded in the PL cortex, a meaningful subset (approximately 35%) of pyramidal neurons depolarized in response to SST, suggesting that polysynaptic SSTR effects may lead to G_i/o_-mediated disinhibitory effects in a subset of pyramidal neurons. To determine whether the hyperpolarizing effects of SST are dependent on polysynaptic activity, action potentials were blocked and monosynaptic circuits isolated with 500 nM TTX. When polysynaptic network activity was blocked, all neurons hyperpolarized in response to SST. SST significantly hyperpolarized pyramidal neurons in both and females (*t_4_*= 3.448, *p = 0.0261*) and males (*t_6_* = 5.095, *p = 0.0022*; **Figure 1K** overall effect separated by sex, with no clear sex differences). This suggests that SST effects on pyramidal neurons are strongest when isolated from polysynaptic circuits, and that the paradoxical depolarization seen in a subset of pyramidal neurons is driven by changes in local network activity. To further confirm that SST is acting to hyperpolarize pyramidal cells independent of both network activity and synaptic (glutamate and GABA) influence, both action potentials and synaptic signaling were blocked using 500 nM TTX, 25µM Picrotoxin to block GABA_A_ receptors, 1µM CGP 55845 to block GABA_B_ receptors, and 3 mM Kynurenic acid to block AMPA and NMDA receptors (**Figure 1L**). SST significantly hyperpolarized pyramidal neurons in both females and males (females *t_6_*= 6.850, *p = 0.0005*; males *t_6_* = 3.179, *p = 0.0191*) when action potentials and synaptic signaling were blocked. This overall demonstrates that SST hyperpolarizes pyramidal cells, through SST receptors on pyramidal cells, independent of changes in synaptic activity. Therefore, the range of SST modulation of pyramidal cells is dependent on the net effect of both postsynaptic hyperpolarization, as well as presynaptic changes in neighboring connections.

Our group has demonstrated we can reliably evoke SST release following 10 Hz optogenetic activation of SST cells in the PL^22^. We next sought to uncover whether *endogenously*-evoked SST release exerted a similar effect on PL layer 2/3 pyramidal cells. To combine this method with our whole-cell electrophysiological approach, we optogenetically activated ChR2-expressing SST cells at 10 Hz in the presence of GABA receptor blockers (25 µM Picrotoxin to block GABA_A_ receptors and 1 µM CGP 55845 to block GABA_B_ receptors) to exclude the effect of co-released GABA, while simultaneously recording from pyramidal neurons (**Figure 1N and 1O**). SST cells fired action potentials in response to a brief 10 Hz (470 nm) stimulation comparable to our previously published work (**Figure 1P**). To assess peptide release, SST cells were optogenetically stimulated at 10Hz (470 nm) for 10 min while simultaneously recording the membrane potential of pyramidal cells (GABA receptor antagonists included in the aCSF; **Figure 1Q**). Males and females were visualized separately to assess whether *endogenously*-evoked SST differed across sexes. 10 min of 10 Hz stimulation did not significantly alter the membrane potential in females (*t_6_* = 0.02415, *p = 0.9815*) or males (*t_11_* = 1.801, *p = 0.0992*; **Figure 1R** overall effect separated by sex). Further, when 1 µM of the SST receptor antagonist cyclo-SST was added to the bath to block SST receptors along with synaptic blockers, 10 min of 10Hz stimulation did not significantly alter the membrane potential in females (*t_6_* = 0.09895, *p = 0.9244*) or males (*t_4_* = 2.263, *p = 0.0864*; **Figure 1S** overall effect separated by sex). Optogenetic stimulation of SST neurons produced a similarly variable effect on membrane potential as that seen with SST bath application when network activity was not blocked (**Figure 1J**). There was also greater standard deviation observed in the aCSF group (1.770; **Figure 1 Q)** than when SSTRs were blocked (1.383; **Figure 1 R**) suggesting SST may contribute to this variability. Therefore, data from **Figure 1R** were split into thirds for visualization to mimic the effect seen with bath application (as approximately 1/3 of cells hyperpolarized, 1/3 remained stable, and 1/3 depolarized in response to SST; **Figure 1T**). There was no effect of optogenetic stimulation of SST neurons in the presence of the SSTR antagonist cyclo-SST (*t_11_* = 1.381, *p = 0.1946* **Figure 1U**). Interestingly, there was a visible depolarization observed in the presence of cyclo-SST, which implicates co-release of other neuropeptides beyond SST. Collectively, these experiments demonstrate the hyperpolarizing effect of SST on pyramidal neurons under various experimental conditions and reveal the complexity of SST peptidergic function. These experiments led us to further examine the mechanisms by which SST modulates pyramidal neurons and the (if any) reversibility of this effect, the effect of the SST on other neurons with the PL, to more deeply interrogate the conditions under which it may be released, and to explore effects of this peptide on behavior.

#### SST actions are blocked, but not reversed, by SSTR antagonists

Next, to confirm that exogenous SST bath application effects, and opto-stimulation effects, were mediated specifically by SST receptors, we performed slice electrophysiology experiments in the presence of a broad SSTR antagonist. 1 µM cyclo-SST was included in the aCSF prior to and throughout application of SST. When cyclo-SST was present in the aCSF, SST did not significantly alter the RMP (representative circuit diagram in **Supplemental Figure 1A**, rheobase traces in **Supplemental Figure 1B**, bath application in **Supplemental Figure 1C,** and summary RMP in **Supplemental Figure 1D**). SST had no effect on rheobase at RMP (**Supplemental Figure 1E**) or action potential threshold at RMP (**Supplemental Figure 1F**). Further, SST had no effect on the rheobase at –70 mV (**Supplemental Figure 1G**) or action potential threshold at –70 mV (**Supplemental Figure 1H**). Moreover, SST did not affect the number of action potentials fired in the VI plot at either RMP or –70 mV (**Supplemental Figure 1I-L**). Together, these experiments confirm that SST-mediated hyperpolarization and reduced excitability of pyramidal neurons is dependent on SSTRs. This experiment also serves as an additional control and further validating results observed in **Figure 1** and demonstrating cell stability in the presence of the antagonist throughout the long time-course of our experiments.

We further determined whether SST-mediated hyperpolarization and reduced excitability of pyramidal neurons are reversible with post-application of an SSTR antagonist. Pyramidal neurons in layer 2/3 of the PL cortex were patched and 1 µM SST applied followed by 1 µM cyclo-SST application. Results were comparable to those without post-application of the antagonist in **Figure 1**. SST with a cyclo-SST washout still significantly hyperpolarized the RMP and rheobase at RMP (representative circuit diagram in **Supplemental Figure 2A**, rheobase traces in **Supplemental Figure 2B**, overall bath application of SST and cyclo-SST washout in **Supplemental Figure 2C**, and overall RMP effect in **Supplemental Figure 2D**). While the rheobase at RMP was significantly increased (**Supplemental Figure 2E**), the action potential threshold at RMP, rheobase at – 70 mV, and action potential threshold at –70 mV were not significantly changed from pre-SST baseline (**Supplemental Figure 2F-H**). This protocol resulted in no significant change in the number of action potentials fired in the VI plot at both RMP and –70 mV (**Supplemental Figure 2I-2L**). This suggests that overall, the effect of SSTR activation on RMP and rheobase is largely not reversible with an SSTR antagonist.

**Figure 2.**
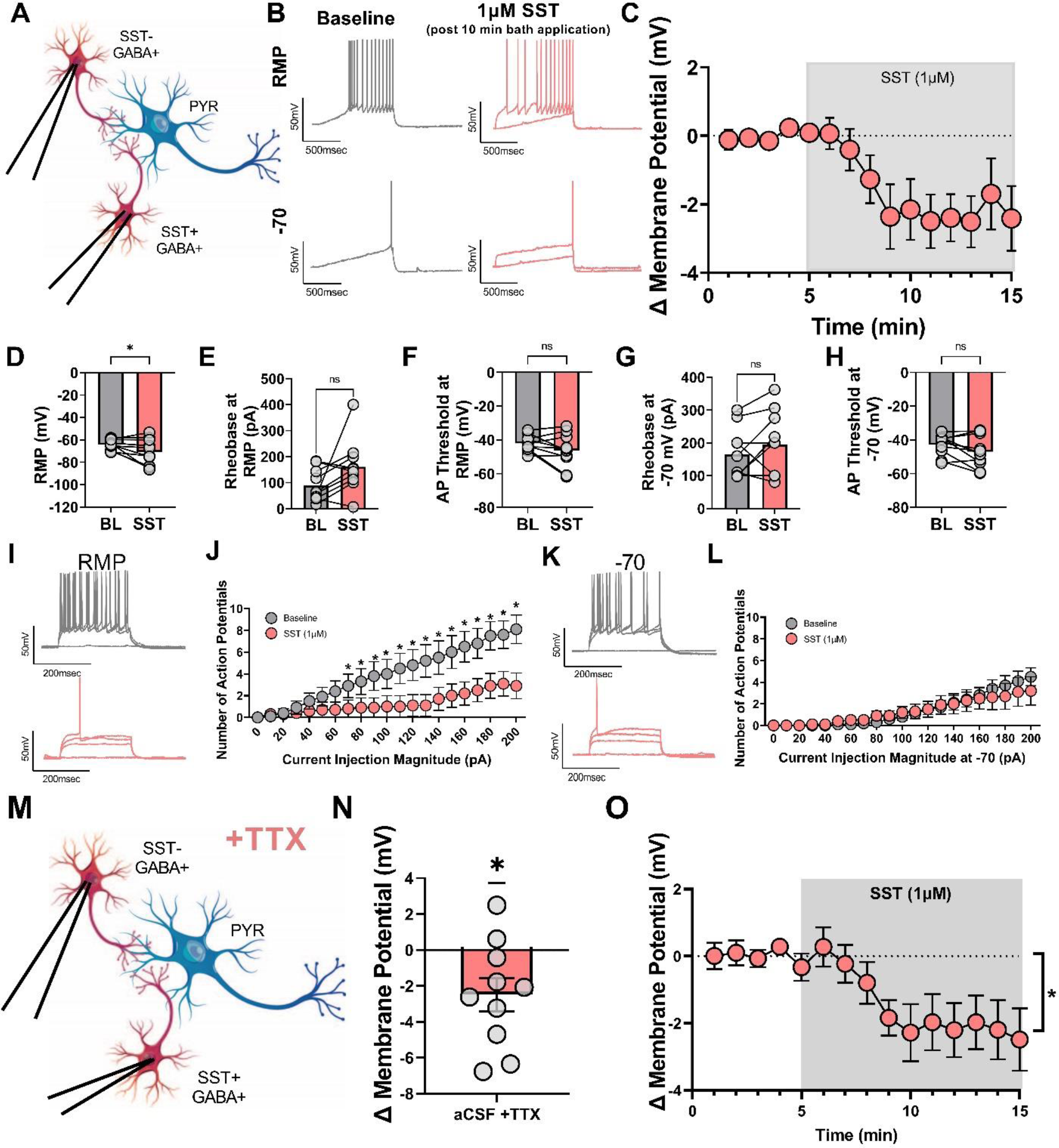
SST dampens excitability of and hyperpolarizes PL non-pyramidal cells. **(A)** Schematic of experimental setup. Whole-cell current clamp recordings were conducted in PL cortex layer 2/3 GABA neurons. **(B)** Representative traces before (grey) and after (red) SST 1 µM application at both RMP (top) and –70 mV (bottom) for rheobase experiments. **(C)** Change in membrane potential over time following SST bath application**. (D)** SST significantly decreased the RMP. However, **(E)** the rheobase at RMP, and **(F)** the action potential threshold at RMP were not significantly changed**. (G)** In addition, the rheobase at –70 mV, and **(H)** the action potential threshold at –70 mV were not significantly altered. **(I)** Representative VI traces at RMP (corresponding to 0, 110, 150, 190 pA of injected current) before (grey) and after (red) SST application. **(J)** SST significantly reduces the number of action potentials fired in response to increasing amounts of current injection at RMP. **(K)** Representative VI traces at –70 mV (corresponding to 0, 110, 150, 190 pA of injected current) before (grey) and after (red) SST application. **(L)** SST does not significantly alter the number of action potentials fired at the common holding potential of –70 mV. **(M)** Model depicting isolation of network independent effects using TTX while patching PL cortex non-pyramidal neurons. **(N)** Change in membrane potential over time following SST bath application. **(O)** Time-course of SST effects. For panels A-L, n= 10 cells from 2 female and 4 male mice. For panels M-O, n=10 cells from 4 female and 4 male mice.

#### SST hyperpolarizes and decreases excitability of PL cortical GABAergic neurons

After demonstrating the effect of SST on glutamatergic pyramidal neurons in the PL in **Figure 1**, we sought to elucidate the effect of SST on the other major population of neurons in the PL, GABAergic neurons, as network-intact experiments clearly implicated polysynaptic regulation of pyramidal neurons by SST. *Ex vivo* whole-cell current clamp recordings (identical to **Figure 1**) were conducted in non-pyramidal, GABAergic populations in the PL cortex (representative circuit diagram in **Figure 2A**, distinguished by morphology, membrane characteristics, and electrophysiological properties as outlined above^3^). Representative traces of rheobase recordings before and after SST application are shown in **Figure 2B**. SST hyperpolarized non-pyramidal neurons and significantly decreased the RMP (*t*_9_ = 2.291, *p* = 0.0477; **Figure 2C-D**). SST did not significantly alter the rheobase at RMP (*t*_9_ = 2.046, *p* = 0.0711; **Figure 2E**), action potential threshold at RMP (*t*_9_ = 1.751, *p* = 0.1138; **Figure 2F**), rheobase at –70 mV (*t*_9_ = 1.320, *p* = 0.2195; **Figure 2G**), or action potential threshold at –70 mV (*t*_9_ = 1.840, *p* = 0.0990; **Figure 2H**). However, SST significantly reduced the number of action potentials fired in response to increasing current injection at RMP (2-way ANOVA; *F*_current_(20,180) = 22.72, *p* <0.0001; *F*_drug_(1,9) = 6.198, *p* = 0.344, *F*_current x drug_ (20,180) = 8.437, *p* <0.0001, **Figure 2I-J;** significant post-hoc Bonferroni’s are indicated on figure) but not at –70 mV (2-way ANOVA; *F*_current_(20,180) = 14, p <0.0001; *F*_drug_(1,9) = 0.0183, *p* = 0.8951, *F*_current x drug_ (20,180) = 1.0, *p* = 0.4063; **Figure 2K-L**). These findings suggest that SST also acts on GABAergic populations within the PL cortex. The inter-spike interval (ISI) of GABA cell action potentials did not significantly correlate with the magnitude of SST modulation (change in membrane potential (mV); r(10) = 0.07, *p* = 0.847; data not shown) suggesting no clear differences in SST effects on putatively fast-spiking GABA neurons versus non-fast spiking GABA neurons. When direct, network-independent effects (representative circuit diagram in **Figure 2M**) were isolated using TTX (500 nM), SST significantly hyperpolarized non-pyramidal cells (*t_9_* = 2.668, *p* = 0.0257; **Figure 2N** for individual data and **Figure 2O** for time course data).

#### SST neurons display task-relevant activity during exploratory behaviors

### Overall behavior

After establishing the pharmacological effect of SST in the PL cortex, we sought to determine the behavioral conditions under which this population of SST cells become activated. To do this, we explored whether *in vivo* activity of SST neurons was related to behavioral performance in the EPM and OFT. VIP::Cre;SST::Flp mice were injected in PL cortex with dual AAVs encoding a Flp-dependent GCaMP6f and Cre-dependent TdTomato, and implanted with an optical fiber in PL cortex (schematic and representative histology, **Figure 3A-B**; histology from all mice in **Supplemental Figure 3A**). A custom-made dual-color spectrometer-based *in vivo* fiber photometry system was used (schematic in **Figure 3C**, sample fluorescence spectrum in **Figure 3D**). No sex differences were detected in the analyzed behavior or photometry signals, so data were pooled by sex (**Figure 3** and **Supplemental Figure 3**; pink dots denote females, purple dots denote males).

**Figure 3.**
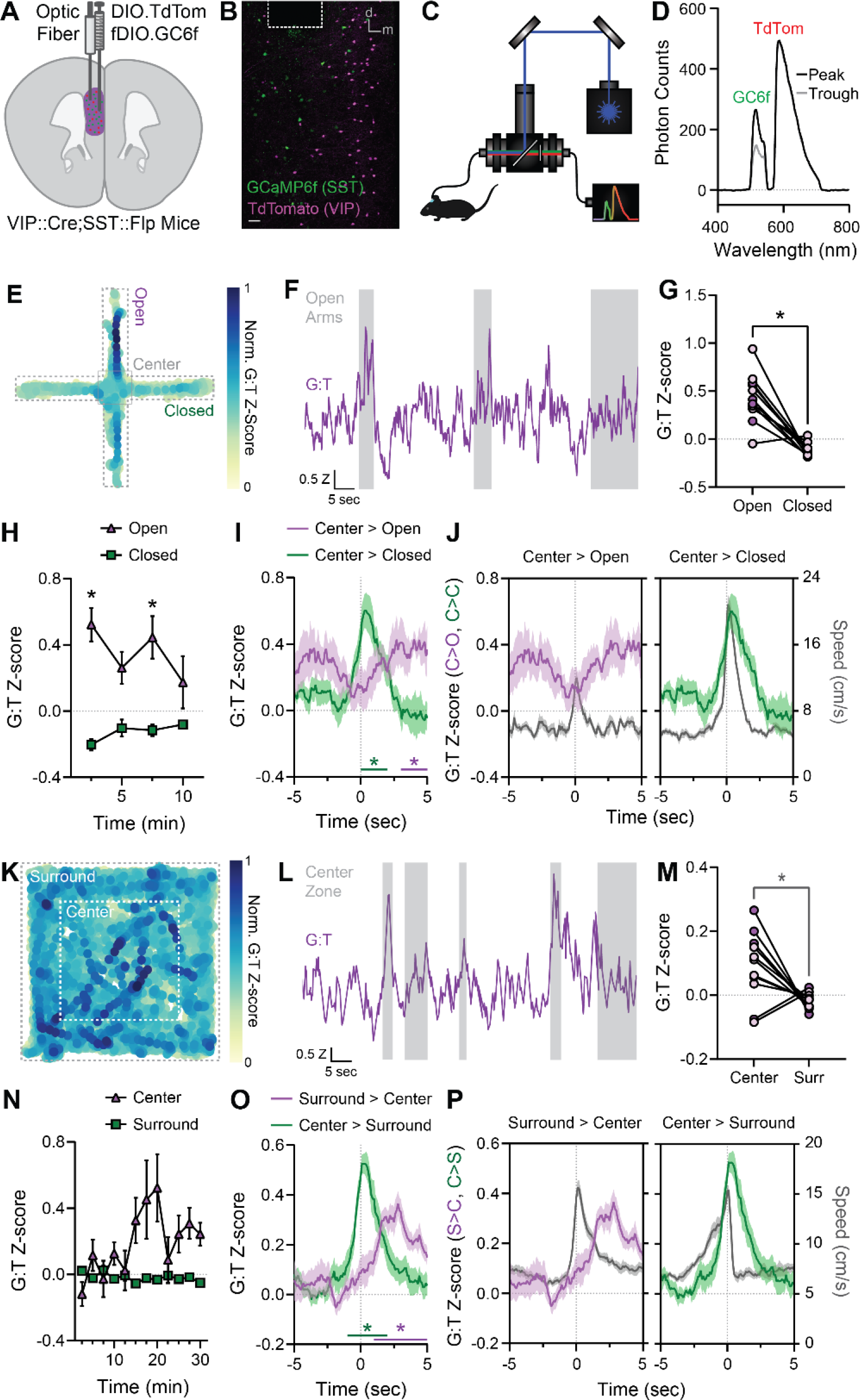
Somatostatin neurons in PL are active during exploration of the EPM and OFT. **(A)** Schematic of AAV9.CAG.DIO.TdTomato and AAV9.Ef1a.fDIO.GCaMP6f injection with optic fiber implant into unilateral PL cortex of VIP::Cre;SST::Flp mice. **(B)** Sample image of GCaMP6f and TdTomato expression (in putative SST+ and VIP+ interneurons, respectively) and optic fiber placement in PL cortex. Scale bar: 50 µm. **(C)** Schematic of dual-color spectrometer-based *in vivo* fiber photometry system. **(D)** Sample fluorescence spectrum from PL cortex showing putative GCaMP6f and TdTomato fluorescence at a peak and trough of the GCaMP6f response. **(E)** Sample heatmap of PrL SST+ interneuron Ca^2+^ dynamics as a function of location in the EPM. Ca^2+^-dependent GCaMP6f fluorescence intensity is reported as the z-scored ratio of GCaMP6f:TdTomato (G:T) fluorescence, normalized between 0 and 1. **(F)** Sample photometric recording of G:T fluorescence during EPM exploration. Grey bars mark periods of open arm exploration. **(G)** Mean z-scored G:T fluorescence during exploration of the open vs. closed arms of the EPM. Dots represent individual mice (N = 11). Male mice, purple; female mice, pink. **(H)** Mean z-scored G:T fluorescence during exploration of the open vs. closed arms across the 10-min EPM test. **(I)** Z-scored G:T fluorescence aligned to center-to-open arm and center-to-closed arm transitions in the EPM, averaged across mice (N=11). **(J)** Z-scored G:T fluorescence and mouse speed aligned to center-to-open arm (left) and center-to-closed arm (right) transitions in the EPM, averaged across mice (N = 11). **(K)** Sample heatmap of z-scored G:T fluorescence (normalized between 0 and 1) as a function of location in the OF. **(L)** Sample photometric recording of G:T fluorescence during OF exploration. Grey bars mark periods of center zone exploration. **(M)** Mean z-scored G:T fluorescence during exploration of the center and surround of the OF. Dots represent individual mice (N = 11). Male mice, purple; female mice, pink. **(N)** Mean z-scored G:T fluorescence during exploration of the center and surround across the 30-min OF test. **(O)** Z-scored G:T fluorescence aligned to surround-to-center zone and center-to-surround zone transitions in the OF, averaged across mice (N=11). **(P)** Z-scored G:T fluorescence and mouse speed aligned to surround-to-center zone (left) and center-to-surround zone (right) transitions in the OF, averaged across mice (N=11).

### Elevated Plus Maze

SST neurons showed task-related increases in calcium (Ca^2+^) activity in the EPM (representative heat map of PL SST neuron Ca^2+^ dynamics as a function of location in the EPM in **Figure 3E;** sample photometric recording of GCaMP6f:TdTomato, indicated as G:T, fluorescence during EPM exploration in **Figure 3F**; grey bars denote periods of open arm exploration). On average, SST neuron activity was higher in the open versus closed arms (*t*_10_ = 6.101 *p* < 0.001; **Figure 3G**). This heightened open arm-related activity gradually diminished across the 10-min EPM test (*F*_Time x Arm_(2.87, 22.93) = 3.487, *p* < 0.05; **Figure 3H**). In addition, SST neuron activity was dynamically altered around arm transitions in the EPM. SST neuron activity was generally elevated in the 10 seconds around both transitions from the center of the maze to the open arms, and from the center to the closed arms (**Figure 3I**; one-sample t-tests vs. 0: center>open: *t*_10_ = 4.351, *p* = 0.0014; center>closed: *t*_10_ = 2.861, *p* = 0.0169). SST neuron activity was consistently elevated across transitions from the center to the open arms, with the greatest activity 3-5 seconds after the transition (Bonferroni-corrected post-hoc one-sample t-tests, *: 3-4 sec and 4-5 sec, *p* < 0.05; **Figure 3I**). In contrast, during transitions from the center to the closed arms, SST neuron activity peaked more discretely in the two seconds following transitions f (Bonferroni-corrected post-hoc one-sample t-tests, *: 0-1 sec and 1-2 sec, *p* < 0.05; **Figure 3I**). In line with these findings, SST neuron activity dynamics differed significantly between the two transition types (2-way RM ANOVA: *F*_Time x Transition_(9,180) = 5.672, *p* < 0.0001; significant post-hoc Šidák’s tests: C>C diff. from C>O at 0-1, 3-4, and 4-5 sec, *p* < 0.05).While SST neuronal activity aligned with speed during closed arm transitions, this relationship was not present during open arm transitions (**Figure 3J** **and Supplemental Figure 6**).

### Open Field Test

Task-relevant activity of SST neurons was also seen in the OFT (representative heat map of PL SST Ca^2+^ dynamics as a function of location in the OFT in **Figure 3K**, and sample photometric recording of G:T fluorescence during OFT in **Figure 3L**; grey bars denote periods of center zone exploration). Similar to recordings during the EPM, average SST neuron activity was higher during exploration of the center of the OF relative to the surround zone (*t*_10_ = 2.797, *p* = 0.0189; **Figure 3M**). Unlike the EPM, heightened center zone-related activity progressively developed over the 30-min OFT (*F*_TimexZone_(4.55, 43.39) = 2.759, *p* < 0.05; **Figure 3N**). SST neuron activity was dynamically altered around zone transitions in the OFT. SST neuron activity was generally elevated in the 10 seconds around transitions from the surround to the center, and from the center to surround (**Figure 3O**; one-sample t-tests vs. 0: surround>center: *t*_10_ = 5.069, *p* = 0.0005; center>surround: *t*_10_ = 5.234, *p* = 0.0004). SST neuron activity gradually increased following transitions from the surround to the center zone (Bonferroni-corrected post-hoc one-sample t-tests, *: 1-2 sec, 2-3 sec, 3-4 sec, and 4-5 sec, *p* < 0.005; **Figure 3O**). In contrast, SST neuron activity was particularly high immediately preceding and following transitions from the center to the surround zone (*:-1-0 sec, 0-1 sec, and 1-2 sec, *p* < 0.05; **Figure 3O**). Consistent with these results, SST neuron activity dynamics differed significantly between the two transition types (2-way RM ANOVA: F_Time x Transition_(9,180) = 10.12, *p* < 0.0001; significant post-hoc Šidák’s tests: C>S diff. from S>C at -1-0 sec and 0-1 sec, *p* < 0.05). SST neuron activity during both transition types aligned comparably to mouse speed during transitions (**Figure 3P**). Together, these experiments highlight that SST neurons are engaged during PFC-dependent exploratory behaviors.

#### Administration of the SSTR agonist Octreotide reveals task- and sex-specific effects on exploratory behavior

### Overall behavior

To assess causal effects of SST peptide signaling in the PL cortex during these same exploratory behaviors, the SSTR agonist Octreotide or aCSF control was administered directly to the PL cortex 10 min prior to the OFT or EPM via bilateral cannulas (histology from mice receiving Octreotide or aCSF in **Figure 4A**). Due to the paucity of literature on SST peptidergic effects on cortically-governed behaviors, males and females were analyzed separately.

**Figure 4.**
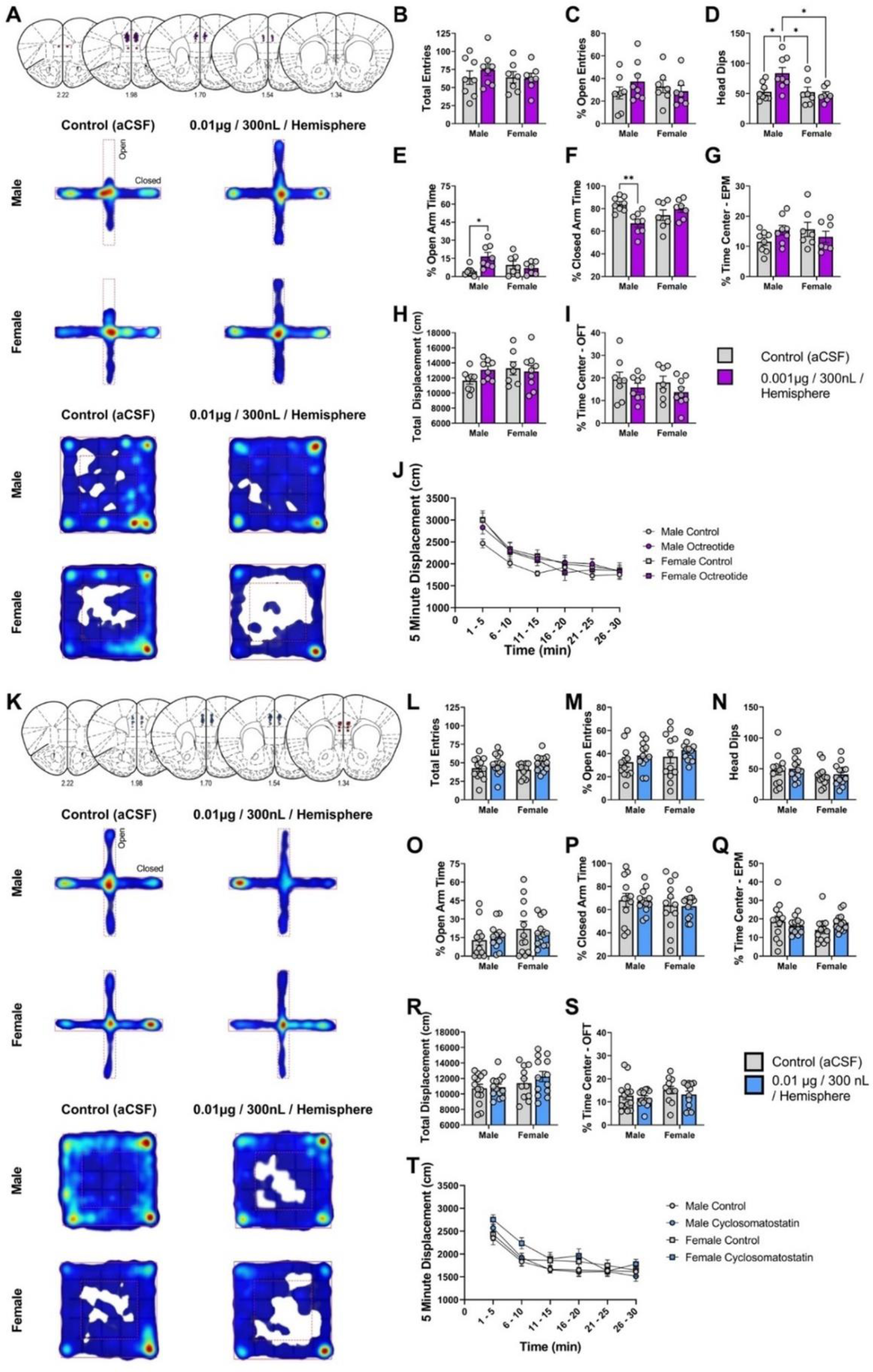
–The SSTR-targeting compound Octreotide in PL decreases avoidance behavior in the EPM, while the SSTR antagonist cyclo-SST has no effect. **(A)** Representative histology and heatmaps for Octreotide behavior. The open arms are oriented vertically, and closed arms are oriented horizontally for all EPM heatmaps. **(B)** Total entries into both open and closed EPM arms. **(C)** Number of entries into the open arm as a percentage of all entries. **(D)** Number of head dip extensions beyond the bounds of the open arm. Octreotide significantly increased the number of head dips in males relative to all other groups. **(E)** Percent time spent in the open arm with respect to total trial time. Octreotide significantly increased the percent of time spent in the open arm for males relative to control. **(F)** Percent time spent in the closed arm with respect to total trial time. Male mice treated with Octreotide significantly decreased the percent of time spent in the closed arm with respect to total trial time. **(G)** Percent time spent in the center zone of the EPM with respect to total trial time. **(H)** Total displacement for mice during the OFT. **(I)** Percent time spent in the center zone of the OFT with respect to total trial time. **(J)** Total combined displacement tracked in five-minute intervals throughout the OFT. **(K)** Representative histology and heatmaps for cyclosomatostatin behavior. The open arms are oriented vertically, and closed arms are oriented horizontally for all EPM heatmaps. **(L)** Total entries into both open and closed EPM arms. **(M)** Number of entries into the open arm as a percentage of all entries. **(N)** Number of head dip extensions beyond the bounds of the open arm. **(O)** Percent time spent in the open arm with respect to total trial time. **(P)** Percent time spent in the closed arm with respect to total trial time. **(Q)** Percent time spent in the center zone of the EPM with respect to total trial time. **(R)** Total displacement for mice during the OFT. **(S)** Percent time spent in the center zone of the OFT with respect to total trial time. **(T)** Total combined displacement tracked in five-minute intervals throughout the OFT. For panels B – G n=30 (8 male control, 8 male Octreotide, and 7 female control, 7 female Octreotide) mice; for panels H – J n=32 (8 male control, 8 male Octreotide,and 7 female control, 9 female Octreodide) mice. For panels L – Q n= 49 (12 male control, 12 male cyclosomatostatin, and 12 female control, 13 female cyclosomatostatin) mice; for panels R – T n = 51 (14 male control, 13 male cyclosomatostatin, and 11 female control, 13 female cyclosomatostatin) mice.

### Elevated Plus Maze

Local administration of the SSTR targeting compound Octreotide (0.001 µg / 300nL / hemisphere) in the PL cortex had no effect on total arm entries or percent open arm entries (representative heat maps in **Figure 4A**; 2-way ANOVA; *F_sex_*(1,26) = 0.4680, *p =* 0.5000; *F*_drug_(1,26) = 0.3289, *p* = 0.5712, *F*_sex x drug_ (1,26) = 0.5580, *p* =0.4618; **Figure 4B**, significant post-hoc Tukey’s are indicated on figure; 2-way ANOVA; *F_sex_*(1,26) = 0.06454, *p =* 0.8015; *F*_drug_(1,26) = 0.2855, *p* = 0.5976, *F*_sex x drug_ (1,26) = 1.598, *p* =0.2174; **Figure 4C**, significant post-hoc Tukey’s are indicated on figure). Interestingly, Octreotide-treated male mice, but not female mice, showed a significant increase in the number of head dips over the open arm of the EPM (2-way ANOVA; *F_sex_*(1,26) = 5.917, *p=0.0222*; *F*_drug_(1,26) = 3.264, *p* = 0.0824, *F*_sex x drug_ (1,26) = 5.614, *p* =0.0255; **Figure 4D**; significant post-hoc Tukey’s are indicated on figure). There was also a significant interaction between drug and sex for percent time in the open and closed arms (2-way ANOVA; *F_sex_*(1,26) = 0.6452, *p =* 0.4291; *F*_drug_(1,26) = 3.462, *p* = 0.0741, *F*_sex x drug_ (1,26) = 7.868, *p* =0.0094; **Figure 4E**, significant post-hoc Tukey’s are indicated on figure; 2-way ANOVA; *F_sex_*(1,26) = 0.1647, *p =* 0.6882; *F*_drug_(1,26) = 2.868, *p* = 0.1023, *F*_sex x drug_ (1,26) = 10.75, *p* =0.0030; **Figure 4F**; significant post-hoc Tukey’s are indicated on figure), with a significant increase and decrease, respectively, seen only in male mice treated with Octreotide relative to control. There was also no significant effect of Octreotide administration on the percent of time spent in the center of the EPM (2-way ANOVA; *F_sex_*(1,26) = 0.2568, *p=0.6166*; *F*_drug_(1,26) = 0.1419, *p* = 0.7095, *F*_sex x drug_ (1,26) = 3.117, *p* =0.0892; **Figure 4F**; significant post-hoc Tukey’s are indicated on figure). This suggests that administration of an SSTR agonist alters exploratory behavior in a novel context in male, but not female, mice.

### Open Field Test

Octreotide administered into PL cortex had no effect on the overall distance traveled in the OFT (2-way ANOVA; *F*_sex_(1,28) = 1.054, *p=0.3134*; *F*_drug_(1,28) = 0.5550, *p* = 0.4625, *F*_sex x drug_ (1,28) = 1.855, *p* = 0.1840; representative heat maps in **Figure 4A**; distance traveled in **Figure 4H**), further suggesting the changes seen in the EPM were not due to alterations in gross motor behavior. In addition, Octreotide did not alter the percent time in the center in either sex (2-way ANOVA; *F_sex_*(1,28) = 0.4335, *p=0.5147*; *F*_drug_(1,28) = 2.426, *p* = 0.1306, *F*_sex x drug_ (1,28) = 0.02013, *p* =0.8882; **Figure 4I**). Octreotide did not significantly alter displacement when time was binned by 5 min (3-way ANOVA; *F_time_*(3.716, 104.0) = 67.10, *p<0.0001*, *F_sex_*(1, 28) = 1.054, *p = 0.3134*, *F_drug_*(1, 28) = 0.5550, *p = 0.4625*, *F_time_* _x sex_(5, 140) = 2.638, *p = 0.0260*, *F_time x drug_*(5, 140) = 0.7609, *p = 0.5795*, *F_sex x drug_*(1, 28) = 1.855, *p = 0.1840*, *F_time x sex x drug_*(5, 140) = 0.4731, *p = 0.7959*; **Figure 4J**). This suggests that the changes in exploratory behavior induced by Octreotide were unique to the EPM and not driven by changes in general ambulatory behavior.

#### Administration of the SSTR antagonist cyclosomatostatin does not alter ambulatory or exploratory behavior

### Overall behavior

After establishing a role for exogenous administration of SST targeting compound Octreotide to the PL in promoting exploratory behavior, we sought to determine whether we could uncover the role of endogenous SST during these same exploratory behaviors by blocking SSTRs. The pan-SSTR antagonist cyclo-SST or aCSF control was administered directly to the PL cortex 10 min prior to the OFT or EPM via bilateral cannulas similarly to the experiments conducted with Octreotide (histology from mice receiving cyclo-SST or aCSF in **Figure 4K**). Importantly, while our slice electrophysiology experiments suggested this dose was effective at blocking SST actions, they also show that SSTR effects are non-reversible. Thus, we considered these experiments a non-perfect approach to uncovering endogenous peptide release.

### Elevated Plus Maze

Local administration of the SSTR antagonist cyclo-SST (0.01 µg / 300nL / hemisphere) in the PL cortex had no effect on total arm entries or percent open arm entries (representative heat maps in **Figure 4K**; 2-way ANOVA; *F*_sex_(1,45) = 0.01162, *p = 0.91*46; *F_drug_*(1,45) = 0.3.727, *p = 0.0598*, *F_sex x drug_*(1,45) = 0.2008, *p =0.6562*; **Figure 4L**, significant post-hoc Tukey’s are indicated on figure; 2-way ANOVA; *F_sex_*(1,45) = 1.353, *p = 0.2509*; *F_drug_*(1,45) = 1.974, *p = 0.1668*, *F_sex x drug_* (1,45) = 1.410e-005, *p = 0.9970*; **Figure 4M**, significant post-hoc Tukey’s are indicated on figure). There was no significant increase in the number of head dips over the open arm of the EPM (2-way ANOVA; *F_sex_*(1,45) = 2.414, *p=0.1273*; *F_drug_*(1,45) = 0.1363, *p = 0.7137*, *F_sex x drug_* (1,45) = 0.002505, *p = 0.9603*; **Figure 4N**; significant post-hoc Tukey’s are indicated on figure). There was also no significant interaction between drug and sex for percent time in the open and closed arms (2-way ANOVA; *F_sex_*(1,45) = 1.877, *p = 0.1775*; *F_drug_*(1,45) = 0.0005470, *p = 0.9814*, *F_sex x drug_* (1,45) = 0.6088, *p =0.4393*; **Figure 4O**, significant post-hoc Tukey’s are indicated on figure; 2-way ANOVA; *F_sex_*(1,45) = 0.8560, *p = 0.3598*; *F_drug_*_(_1,45) =0.03603, *p = 0.8503*, *F_sex x drug_*(1,45) = 0.0009172, *p = 0.9760*; **Figure 4P**; significant post-hoc Tukey’s are indicated on figure). There was no significant effect of cyclo-SST administration on the percent of time spent in the center of the EPM (2-way ANOVA; *F_sex_*(1,45) = 0.5035, *p=0.4816*; *F_drug_*(1,45) = 0.1820, *p = 0.6717*, *F_sex x drug_* (1,45) = 2.567, *p = 0.1161*; **Figure 4Q**; significant post-hoc Tukey’s are indicated on figure). This demonstrates that administration of a SSTR antagonist has no effect on exploratory or ambulatory behavior in a novel context in male or female mice.

### Open Field Test

Cyclo-SST administered into PL cortex had no effect on the overall distance traveled in the OFT (2-way ANOVA; *F_sex_*(1,47) = 3.300, *p = 0.0757*; *F_drug_*(1,47) = 0.7976, *p = 0.3764*, *F_sex x drug_*(1,47) = 0.4138, *p = 0.5232*; representative heat maps in **Figure 4K;** distance traveled in **Figure 4R**), suggesting the changes seen in the EPM were not due to alterations in gross motor behavior. In addition, cycl0-SST did not alter the percent time in the center in either sex (2-way ANOVA; *F_sex_*(1,47) = 1.773, *p = 0.1894*; *F_drug_*(1,47) = 0.9509, *p = 0.3345*, *F_sex x drug_*(1,47) = 0.08929, *p = 0.7664*; **Figure 4S**). Cyclo-SST did not significantly alter displacement when time was binned by 5 min (3-way ANOVA; *F_time_*(5,235) = 80.45, *p < 0.0001*, *F_sex_*(1,47) = 3.300, *p = 0.0757*, *F_drug_*(1, 47) = 0.7976, *p = 0.3764*, *F_time x sex_*(5,235) = 0.9541, *p = 0.4468*, F_time x drug_(5, 235) = 3.201, *p = 0.0082*, *F_sex x drug_*(1, 47) = 0.4138, *p = 0.5232*, *F_time x sex x drug_*(5,235) = 0.8377, *p = 0.5241*; **Figure 4T**).This suggests SST signaling does not drive changes in general ambulatory behavior.

While no broad effects were seen when blocking SSTRs at the dose of cyclo-SST selected, due to the non-reversible nature of SSTR activation (coupled with likely rapid internalization of the receptor, occluding the ability to antagonize it), we caution against interpreting this as a *lack* of endogenous SST signaling in modulating exploratory behavior.

### Validation of Octreotide in *ex vivo* electrophysiology experiments

Finally, in order to confirm that the SST-like agonist Octreotide has similar *ex vivo* effects on pyramidal neurons to SST, we performed identical whole-cell current clamp experiments to those conducted with SST but using 3.27 µM Octreotide (corresponding to the concentration used for behavior; **Supplemental Figure 4**). Representative traces of rheobase recordings are shown in **Supplemental Figure 4B and 4N** for females and males, respectively. Octreotide had no significant effect on RMP in females (**Supplemental Figure 4D**); however, in males Octreotide significantly reduced RMP (**Supplemental Figure 4P**). Octreotide significantly increased the rheobase at RMP in both females (**Supplemental Figure 4E**) and males (**Supplemental Figure 4Q**). Octreotide did not significantly alter the action potential threshold at RMP in females (**Supplemental Figure 4F**); however, Octreotide did significantly decrease the action potential threshold at RMP in males (**Supplemental Figure 4R**). Octreotide significantly increased the rheobase at –70 mV in females (**Supplemental Figure 4G**); however, there was no significant change in the rheobase at –70 mV in males (**Supplemental Figure 4S**). Octreotide did not significantly change the action potential threshold at – 70 mV in females (**Supplemental Figure 4H**) or males (**Supplemental Figure 4T**). Octreotide significantly decreased the number of action potentials fired in the VI plot at both RMP and –70 mV in both females (**Supplemental Figure 4J, 4L**) and males (**Supplemental Figure 4V, 4X**). Overall, while small differences between the sexes and between SST and Octreotide were seen, collectively these data confirm that Octreotide has similar effects to SST in hyperpolarizing and reducing the excitability of pyramidal neurons.

## DISCUSSION

Here, we provide the first evidence that SST peptide signaling in the PL cortex acts to broadly dampen cortical circuits in both male and female mice using both exogenous electrophysiological models and optogenetically evoked endogenous release (**Figure 1-2**). We found that SST neurons are preferentially activated when mice explore the open arms of the EPM and the central zone of the OFT, and during transitions between these zones (**Figure 3**), suggesting that SST neurons are recruited in task-relevant exploratory behaviors. The pro-exploratory effects of intra-PL SSTR agonist administration (**Figure 4**) aligned with some of the SST neuron activity dynamics recorded with fiber photometry (notably the Octreotide-induced increase in open arm time in the EPM and increased SST neuron activity in the open arms of the EPM), though no clear effects were seen with intra-PL administration of the broad SSTR antagonist cyclo-SST. These findings support the overall hypothesis that increased SST neuronal activity observed during the fiber photometry experiments could correspond to endogenous peptide signaling during exploratory behaviors. Collectively, these data suggest that SST and SSTR-targeting compounds alter behavior through inhibition of PL pyramidal neuron outputs and support the need for further investigation teasing apart peptidergic and neurotransmitter actions in these circuits and behaviors.

### SST signaling dampens PL cortical output circuits

Results from the present study fill a critical gap in the literature by providing the first evidence that SST acts as an independent signaling molecule to regulate PL cortical neurons. This work demonstrates that SST reduces membrane potential and intrinsic excitability of both glutamatergic (pyramidal) projection populations and GABAergic (non-pyramidal) local microcircuit neurons in the PL cortex (**Figures 1-2**). We found that SST in the PL cortex acts via activation of SSTRs (**Supplemental Figure 1**) to modulate activity of output pyramidal neurons through monosynaptic and polysynaptic mechanisms (**Figures 1-2**). While pyramidal neurons broadly hyperpolarized in response to SST, when network activity was not blocked, the response was variable, and a subset (approximately 1/3) depolarized (**Figure 1**). When network activity was blocked, all pyramidal neurons hyperpolarized in response to SST (**Figure 1**). This depolarization was therefore likely due to polysynaptic mechanisms (e.g., polysynaptic GABA neuron-mediated disinhibition of pyramidal neurons). Further, SST-induced hyperpolarization occurs independent of synaptic activity (**Figure 1**). This suggests that SST is acting to hyperpolarize pyramidal neurons directly and likely postsynaptically. While GABAergic neurons also broadly hyperpolarized in response to SST (**Figure 2**) there were some differences observed between pyramidal and GABAergic neurons in their response to SST. Notably, SST did not significantly affect action potential threshold or number of action potentials fired when the cells were held at -70. This may be due to differences in effects on voltage-gated channels. Further, the response of putative GABAergic neurons to SST was more variable when network activity was blocked, indicating that they may have differing expression of SSTRs.

To provide insight into the effect of endogenous SST released from SST cells in the PL, we measured the effect of optogenetically-evoked SST release on pyramidal cell membrane potential in the PL (**Figure 1**). Optogenetic activation (10Hz) of SST cells led to a comparable split in hyperpolarization and depolarization as seen with bath application when network activity is maintained, suggesting optogenetic-stimulation of SST neurons leads to comparable endogenous SST release and subsequent network-driven effects as that seen with exogenous bath application. Interestingly, in the presence of the antagonist cyclo-SST, all hyperpolarization was blocked but we observed depolarization in a large subset of cells. This suggests co-release of another neuropeptide by SST neurons, with the literature pointing to many viable candidates. This experiment highlights the complexity of pharmacological signaling of SST neurons, encourages experiments targeted at the independent effect of SST, and opens the door for future studies to tease apart the neurotransmitters and neuropeptides that contribute to the overall effect of SST cells.

Our results indicate the effect of SST on a given neuron results from the net effect of both pre- and postsynaptic effects, and likely depends on the microcircuit in which it is embedded. An important avenue of future study is to explore the specificity of SST effects on different pyramidal neuron output pathways, whether SST-mediated effects differ across genetically distinct populations of pyramidal and GABAergic neurons, and whether these neurons express differing levels of the various SSTRs. It will be critical to assess whether unique glutamatergic inputs onto SST neurons (e.g., those arising from the hippocampus^6^), drive differing peptidergic effects in PL cortex.

SST-mediated hyperpolarization was not reversed by a SSTR antagonist (**Supplemental Figure 2**). The long-lasting effect of SST^13^, and the non-reversable nature^35^ is consistent with the reported actions of other neuropeptides (e.g., Dynorphin^35^). Moreover, previous work indicates rapid agonist-induced internalization of some subsets of SSTRs^36^. Therefore, it is likely that some SSTRs are internalized after SST administration, limiting antagonist binding. Future work will attempt to elucidate the precise mechanism of this long-lasting and non-reversible effect, as it is key to understanding potential therapeutic strategies that involve SSTR agonism.

Studies have indicated an overall inhibitory role for SST-mediated neuromodulation. SST can inhibit the release of growth hormones from the pituitary^37^, reduce glutamate and GABA transmission onto forebrain cholinergic neurons^38^, and inhibit GABA transmission in both the thalamus^39^ and striatum^40^. SST has been shown to hyperpolarize hippocampal pyramidal cells *in vitro*^41^. However, SST may also have an excitatory response at higher concentrations^42^. SST also reduces excitability in cortical pyramidal neurons in the developing brain^34^. The precise effect of SST likely depends on various factors, such as the particular brain region and cell type probed, the relative expression of SSTRs 1-5, subject age, and the concentration of SST. Prior to this investigation, no studies of SST action had been conducted in the PL cortex, and none bridged its *ex vivo* pharmacological circuit effects with related *in vivo* neuronal signaling and behavioral effects.

Previous work from our group demonstrated that SST peptide is released in PL cortex when SST neurons are optogenetically stimulated at 10 Hz^22^, suggesting that changes in SST neuronal firing rates at relatively low frequencies may correspond to alterations in SST release. Recent rodent studies have implicated PL SST neurons as mediators of various behaviors^16, 17, 43^. The neuromodulatory role of SST in the PL presented in the current study suggests that these prior studies should be interpreted as having potential effects on both GABAergic and peptidergic mechanisms. Moreover, reductions in the number of SST neurons observed following chronic stress^44^ and changes in GABAergic populations in patients with neuropsychiatric diseases^2^ are likely to result in altered SST neuromodulation alongside altered GABAergic function. Such changes in SST modulation may contribute to dysregulation of PL cortical neurons and outputs and ultimately be causal to some disease-relevant behaviors.

Characterizing neuropeptide signaling and neuropeptide release *in vivo* has been particularly challenging for researchers owing to their long-lasting action, diverse pharmacology, and lack-of-effect on traditional ionotropic channels^45^. Previous techniques including microdialysis and fast-scan cyclic voltammetry (FSCV) have provided important insights; however, success has been limited by technological constraints^46^. Importantly, the rapid emergence of new optical biosensors for peptides including SST^47, 48^ will allow for *in vivo* investigation of neurotransmitter versus peptide dynamics. These emerging technologies will allow for the decoupling of SST peptide signaling versus co-released GABA (and potentially other peptides as well). Using these sensors with *in vivo* optical imaging techniques (such as fiber photometry for bulk fluorescence or miniscopes for single-photon imaging) will allow for monitoring of endogenous SST peptide signaling during freely moving behaviors. Future work will investigate the precise nature of SST versus GABA release, including in different behavioral paradigms, in different microcircuits, and the interactions between SST and output-specific pyramidal populations.

### SST neurons in PL are active during exploratory behavior, and SST peptide administration into PL similarly alters exploration

Here, we demonstrate that PL SST neurons broadly signal both zone location and transitions in two separate tasks of exploratory behavior (**Figure 3**). Specifically, our photometry recordings reveal that SST neurons are preferentially activated during exploration of the open versus closed arms of the EPM, and the center versus surround zones of the OF. SST neurons have been previously implicated in exploratory behavior as manipulation of SST neuron activity during the EPM influences open arm behavior^49^. The general increase in SST neuron activity in the open arms of the EPM also corroborates prior work^50^. Notably, SST activity did not consistently correlate with speed, showing a relationship only in transitions to the open arm. Our recordings further reveal that SST neuron activity aligns with discrete transitions, most notably from center-to-closed arms of the EPM and from center-to-surround zones in the OF. Our causal evidence that SSTR signaling in PL cortex promotes EPM open arm exploration aligns well with the heightened activation of PL SST neurons in the open arms of the maze. Heightened activity of SST neurons suggests increased SST release. The release of SST requires repetitive action potentials as SST is stored in dense-core vesicles residing away from the active zone^16^. While *in vivo* imaging strategies covered a broad portion of the PFC, and SST Ca^2+^ dynamics do not directly indicate SST peptide release, this alignment, and our prior reports of SST release in response to sustained low-frequency SST neuron activity^22^, suggests that SST peptide may be released during more sustained periods of open arm exploration. SST may serve as a pro-exploration signal, similar to prior suggestions that SST acts as a “pro-resiliency” peptide^2, 17^. In contrast, SST neuron activation that aligns with rapid transitions from the open arms of the maze may reflect activity that preferentially signals through GABAergic transmission over peptidergic modulation. Our future work will test this hypothesized dichotomy by using, among other approaches, *in vivo* recordings of emerging fluorescent SST sensors.

Importantly, while the fiber photometry experiments did not detect any sex-dependent relationship between SST neuron signaling and exploratory behavior, peptide-induced changes in behavior were only identified in male mice. Increased exploratory behavior after infusion of somatostatin receptor 2 agonists in the hippocampus has been previously reported^51^. Additional studies have suggested behavioral effects of SST following intracerebroventricular infusion^52, 53^ and intra-septal and intra-amygdalar infusion^54^. However, critically, these studies were done exclusively in males, and provided little mechanistic insight. Differences observed in males and females may be due to numerous factors, including potential differences in SSTR density and/or relative prevalence of SSTR subtypes^55^. For example, SSTR density is higher in the human brain of males compared to females which may account for the observed effect in males but not females^55^. SSTR density is also higher in the male rat arcuate nucleus and pituitary^56^. Our current work also did not explore a broad dose range of the SSTR antagonist cyclo-SST. While the concentration chosen was sufficient to block SST-mediated effects in slice electrophysiology experiments, additional studies should be conducted to explore doses *in vivo* – where effects may be uncovered at higher concentrations. Furthermore, future work will explore SSTR subtype expression and density in the PL cortex in male and female mice, and whether biased ligands may provide differing effects.

### Potential pharmacological mechanisms of SST-mediated hyperpolarization

Neuropeptides exhibit diverse effects through the activation of numerous intracellular signal transduction cascades, and rarely exert a single consistent effect across systems. The mechanism of SST-induced hyperpolarization is intricate (involves multiple types of SSTRs, diverse intracellular signaling cascades, and effects on ion channels) and has yet to be fully elucidated. SSTRs are G_i/o_-protein-coupled and inhibit adenylyl cyclase, resulting in decreased intracellular cyclic AMP and intracellular Ca^2+57^. SSTRs can modulate different signal transduction cascades including mitogen-activated protein kinase (MAPK) leading to a multitude of cellular consequences^57^. Prolonged exposure to SST can alter gene expression^58^. One downstream mechanism of action following SSTR activation which was recently characterized in other cortical regions is activation of g-protein coupled inwardly rectifying potassium (GIRK) channels^34, 59, 60^. SST may lead to opening of GIRK channels, increasing potassium (K^+^) conductance and leading to K^+^ efflux - thus hyperpolarizing the cell. This mechanism likely accounts for some of the hyperpolarization observed in electrophysiological recordings (anatomically confined to layer 2/3) in **Figure 1 and 2**. SST can also interact synergistically with neurotransmitter systems such as the dopaminergic system to enhance GIRK activation^61^. Importantly, because there are five known SSTRs, it will be important for future work to tease apart SST effects on a receptor-by-receptor basis. It is also possible that SST modulates neurotransmitter signaling (e.g., increase GABA release or modulating GABA_B_ receptors^62^ as well as interacting with other neuropeptides (e.g., dynorphin or CRF)^63^. Moreover, SST can act both pre- and post-synaptically and has also been shown to modulate neurotransmitter release through both loci^16^. SST in the PL cortex likely acts via numerous mechanisms simultaneously to modulate synaptic plasticity, neurotransmission, and excitability. Pyramidal and GABAergic neurons may respond differently to SST based on various factors (e.g., differences in receptor expression). The specific effect and extent of SST-induced hyperpolarization and decreased excitability is therefore likely to be extremely cell and circuit specific.

### SST as a future therapeutic target

Peptides in the PL cortex have begun to gain increased attention for their role in neuropsychiatric diseases. Clinically, SST in the prefrontal cortex is downregulated in numerous neuropsychiatric disorders such as schizophrenia, bipolar disorder, and major depressive disorder, positioning it as a potential therapeutic target^2^. However, little has been done to understand SST as a signaling molecule and its potential to confer pro-resiliency behaviors. In the present study, we demonstrate a role for SST in modulating the activity of PL cortical pyramidal and GABAergic populations. Further, we show that PL SST neurons are activated during exploratory behavior, and that activating PL SSTRs can promote exploratory behavior. Taken together, these findings suggest that activation of these neurons during behavioral tasks may correspond to endogenous peptide release. Future work should further probe the viability of SSTR-targeting compounds for models relevant to clinical neuropsychiatric disorders.

## CONCLUSIONS

Taken together, our work solidifies SST peptide as a key neuromodulator in the PL cortex, by demonstrating its capacity to dampen excitatory and inhibitory network activity and promote a fundamental aspect of animal behavior.

## ACKNOLWEDGEMENTS

This work was funded by the National Institutes of Health (R01AA209403, R21AA028088, and P50AA017823 to NAC; R01NS078168 and R01NS101353 to PJD; T32NS115667 Training Grant to MSH) and National Institute of Neurological Disorders and Stroke Intramural Research Program (ZIA NS003168). The content of this article is solely the responsibility of the authors and does not necessarily represent the official views of the NIH.

Conceptualization: DFB and NAC. Methodology: DFB, MSH, PJD, DAK, JAG, and NAC. Investigation: DFB, KRG, CMA, TTC, JBM, GCS, NCD, MSH. Visualization: DFB, KRG, CMA, TTC, MSH, DAK, and NAC. Funding acquisition: PJD, DAK, JAG, and NAC. Project administration: PJD, DAK, and NAC. Supervision: PJD, DAK, and NAC. Writing: DFB, PJD, JAG, DAK, and NAC.

---

**Supplemental Figure 1.**
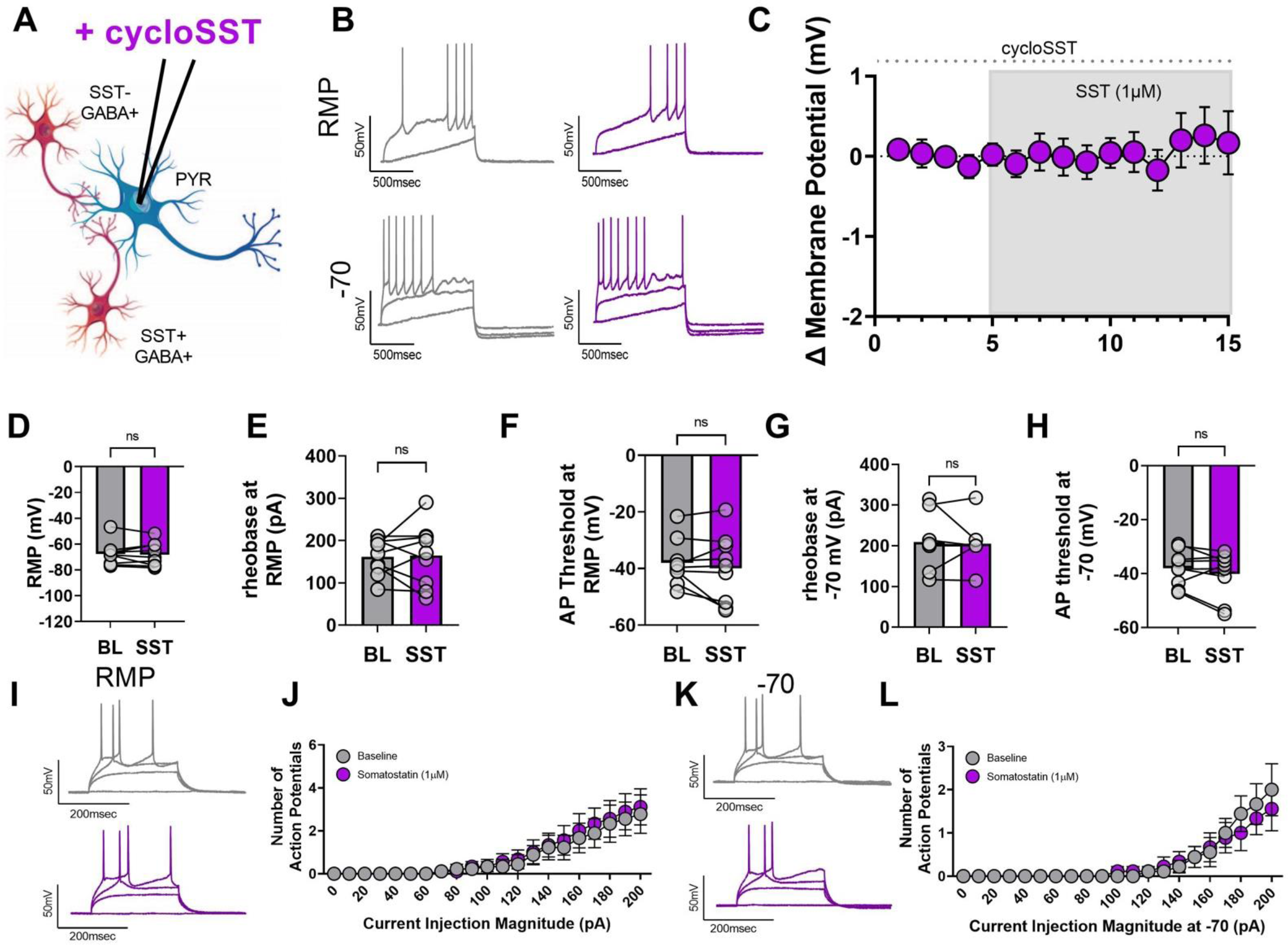
SST-induced alterations in PL pyramidal neuron excitability are dependent on SST receptors. **(A)** Schematic of the experimental setup. Whole-cell current clamp recordings were conducted in PL cortex layer 2/3 pyramidal neurons. **(B)** Representative traces before (grey) and after (purple) 1 µM SST bath application with 1 µM cyclo-SST in the aCSF at both RMP and –70 mV for rheobase experiments. **(C)** Change in membrane potential over time following SST bath application with cyclo-SST 1µM in the aCSF. **(D)** SST does not significantly alter the RMP when 1µM cyclo-SST is present (*t*_8_ = 0.3001, *p* = 0.7717). **(E)** SST did not significantly change the rheobase at RMP in the presence of 1µM cyclo-SST (*t*_8_ = 0.1447, *p* = 0.8885), nor **(F)** the action potential threshold (*t*_8_ = 0.9659, *p* = 0.3624). **(G)** The rheobase at –70 mV (*t*_8_ = 0.3059, *p* = 0.7675), and **(H)** the action potential threshold at –70 mV (*t*_8_ = 1.318, *p* = 0.2240) were not significantly altered. **(I)** Representative VI traces at RMP (corresponding to 0, 110, 150, and 190 pA of injected current). **(J)** SST does not significantly alter the number of action potentials fired in response to increasing amounts of current injection at RMP in the presence of 1µM cyclo-SST (2-way ANOVA; *F*_current_(20,160) = 9.219 *p* <0.0001; *F*_drug_(1,8) = 1.076 *p* = 0.330; *F*_current x drug_ (20,160) = 0.6466 *p* = 0.8721). **(K)** Representative VI traces at –70 mV (corresponding to 0, 110, 150, and 190 pA of injected current). **(L)** SST also had no effect on action potential firing at the standard holding potential of –70 mV in the presence of 1 µM cyclo-SST (2-way ANOVA; *F*_current_(20,160) = 10.24 *p* <0.0001; *F*_drug_(1,8) = 0.1306 *p* = 0.7272; *F*_current x drug_ (20,160) = 1.048 *p* = 0.4104). For all panels, n= 9 cells from 5 female and 3 male mice.

**Supplemental Figure 2.**
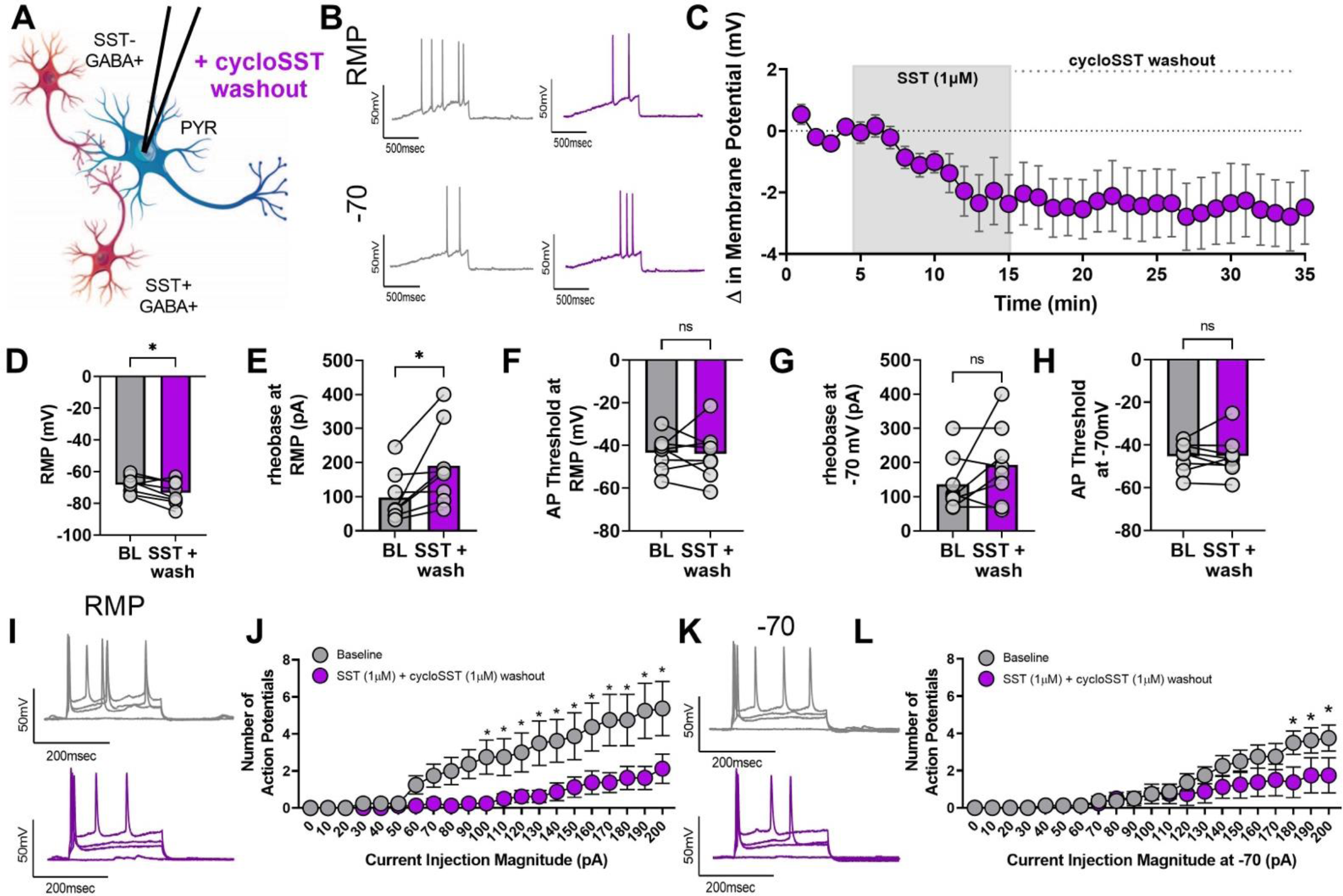
Somatostatin-induced alterations in PL pyramidal neuron excitability are insensitive to reversal by SST antagonists. (A) Schematic of experimental setup. Whole-cell current clamp recordings were conducted in PL cortex layer 2/3 pyramidal neurons. (B) Representative traces before (grey) and after (purple) 1 µM SST followed by a 1 µM cyclo-SST washout at both RMP and –70 mV for rheobase experiments. (C) Change in membrane potential over time following SST bath application and cyclo-SST washout. (D) SST with cyclo-SST washout significantly reduces the RMP (t7 = 2.954, p = 0.0213). (E) While the rheobase at RMP was significantly increased by SST and cyclo-SST washout (t7 = 2.870, p = 0.0240), (F) the action potential threshold was not (t7 = 0.1339, p = 0.8973). (G) The rheobase at –70 mV (t7 = 1.574, p = 0.1595), and (H) the action potential threshold at –70 mV (t7 = 0.1218, p = 0.9065) were not significantly altered. (I) Representative VI traces at RMP (corresponding to 0, 60, 80, and 140 pA of injected current). (J) SST and cyclo-SST washout resulted in a significant interaction between current and SST (2-way ANOVA: Fcurrent(20,140) = 16.20 p <0.0001; Fdrug(1,7) = 4.775 p = 0.0651; Fcurrent x drug (20,140) = 3.170 p <0.0001; significant post-hocs indicated on figure). (K) Representative VI traces at –70 mV (corresponding to 0, 60, 80, and 140 pA of injected current). (L) Number of action potentials fired remained significant reduced at the holding potential of –70 mV (2-way ANOVA; Fcurrent(20,140) = 15.20 p <0.0001; Fdrug(1,7) = 1.595 p = 0.2471; Fcurrent x drug (20,140) = 2.527 p = 0.0009). For all panels, n= 8 cells from 5 female and 2 male mice.

**Supplemental Figure 3.**
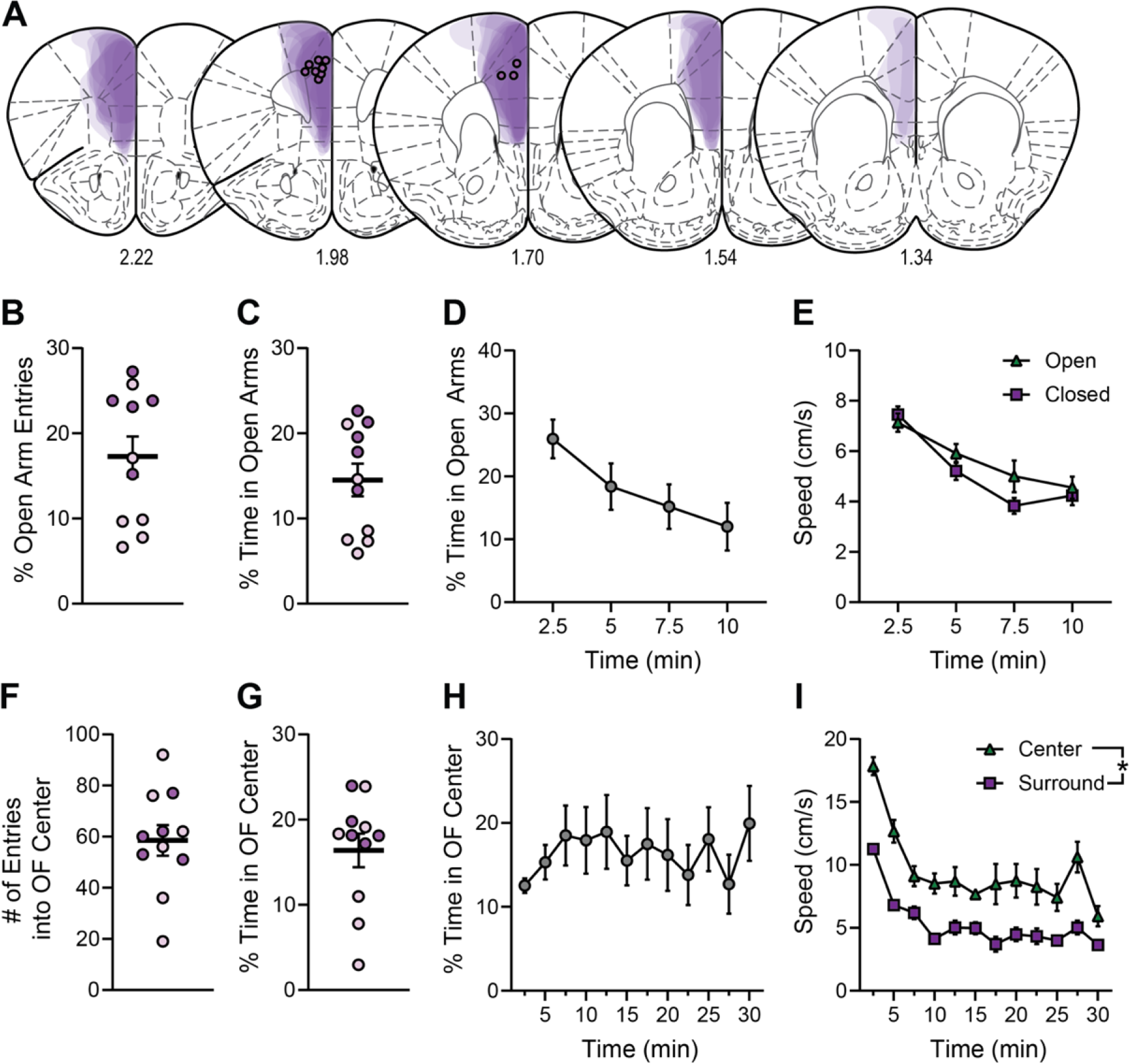
Fiber photometry histology and additional EPM and OFT metrics. (A) Combined GCaMP6f and TdTomato expression (purple) and optic fiber placement in PL cortex. Translucent purple areas denote virus expression in individual mice. (B) Percent number of open arm entries (open/(open+closed) x 100). (C) Percent time spent in open arms (open/(open+closed) x 100). (D) Percent time spent in open arms (open/(open+closed) x 100) across the 10-min EPM test (FTime (2.52, 25.18) = 3.675, p < 0.05). (E) Speed (cm/s) of mice when in the open or closed arms across the 10-min EPM test (2-way RM ANOVA: FTime (2.33,23.25) = 21.17, p < 0.0001). (F) Number of entries made into the center zone of the OFT. (G) Percent time spent in the center zone of the OFT. (H) Percent time spent in the center zone across the 30-min OFT. (I) Speed (cm/s) of mice when in the center or the surround zones across the 30-min OFT (FTime (11,110) = 22.67, p < 0.0001; FZone (1,10) = 70.09, p < 0.0001). Male mice, purple; female mice, pink.

**Supplemental Figure 4.**
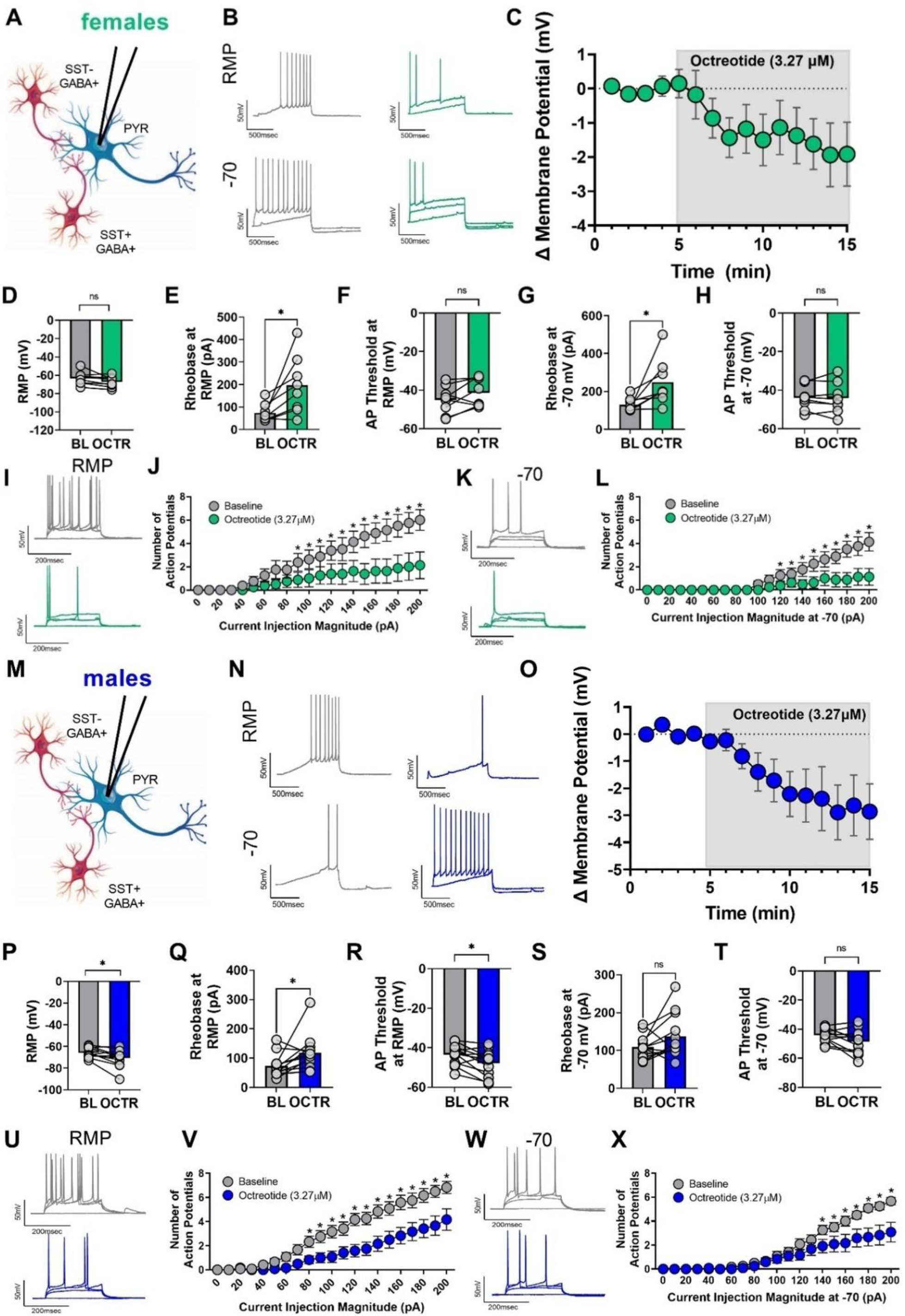
The SST-like agonist Octreotide has similar ex vivo effects to SST on PL neuron physiology in both female and male mice. (A) Schematic of experimental setup. Whole-cell current clamp recordings were conducted in PL cortex layer 2/3 pyramidal neurons in females for panels A-L, and in males for panels M-X. (B) Representative traces before (grey) and after (green) 1 µM SST application at both RMP and –70 mV for rheobase experiments. (C) Change in membrane potential over time following Octreotide bath application. (D) Octreotide did not significantly alter membrane potential (t7 = 2.239 p = 0.0602). (E) The rheobase at RMP was significantly increased (t7 = 2.875, p = 0.0238), and (F) the action potential threshold was not significantly changed (t7 = 1.844, p = 0.1077). (G) In addition, the rheobase at –70 mV was significantly increased (t7 = 2.620, p = 0.0344), and (H) the action potential threshold at –70 mV was not significantly altered (t7 = 0.1792, p = 0.8628). (I) Representative VI traces at RMP (corresponding to 0, 40, 60, 80, 140 pA of injected current) before (grey) and after (green) SST application. (J) Octreotide significantly reduces the number of action potentials fired in response to increasing amounts of current injection at RMP (2-way AVNOA; Fcurrent(20,140) = 13.78; p <0.0001; Fdrug(1,7) = 24.19, p = 0.0017; Fcurrent x drug (20,140) = 11.11, p <0.0001; significant post-hoc Bonferroni’s are indicated on figure). (K) Representative VI traces at –70 mV (corresponding to 0, 40, 60, 80, 140 pA of injected current). (L) Octreotide significantly reduces the number of action potentials fired at the common holding potential of –70 mV (2-way ANOVA; Fcurrent(20,140) = 10.42; p <0.0001, Fdrug(1,7) = 29.27, p = 0.0010, Fcurrent x drug (20,140) = 19.18, p <0.0001; significant post-hoc Bonferroni’s are indicated on figure). (M) Schematic of experimental setup. Whole-cell current clamp recordings were conducted in PL cortex layer 2/3 pyramidal neurons in males for panels M-X. (N) Representative traces before (grey) and after (blue) SST 1 µM application at both RMP and – 70 mV for rheobase experiments. (O) Change in membrane potential over time following Octreotide bath application. (P) Octreotide significantly reduced the RMP in males (t11 = 2.738 p = 0.0193). (Q) The rheobase at RMP was significantly increased (t11 = 2.341 p = 0.0391), and (R) the action potential threshold was significantly reduced (t11 = 2.848 p = 0.0159). (S) In addition, the rheobase at –70 mV was not significantly altered (t11 = 1.745 p = 0.1088), (T) while the action potential threshold at –70 mV was not significantly altered (t11 = 2.200 p = 0.0501). (U) Representative VI traces at RMP (corresponding to 0, 40, 60, 80, 140 pA of injected current) before (grey) and after (green) SST application. (V) Octreotide significantly reduces the number of action potentials fired in response to increasing amounts of current injection at RMP (2-way ANOVA; Fcurrent(20,220) = 75.68; p <0.0001; Fdrug(1,11) = 11.53, p = 0.0060; Fcurrent x drug (20,220) = 5.769, p <0.0001; significant post-hoc Bonferroni’s are indicated on figures). (W) Representative VI traces at –70 mV (corresponding to 0, 40, 60, 80, 140 pA of injected current). (X) Octreotide significantly reduces the number of action potentials fired at the common holding potential of –70 mV (2-way ANOVA; Fcurrent(20,220) = 53.05 p <0.0001; Fdrug(1,11) = 5.129 p = 0.0447; Fcurrent x drug (20,220) = 6.954 p <0.0001; X, significant post-hoc Bonferroni’s are indicated on figures). For panels A-L n = 8 cells from 5 female mice. For panels M-X n = 12 cells from 9 male mice.

**Supplemental Figure 5.**
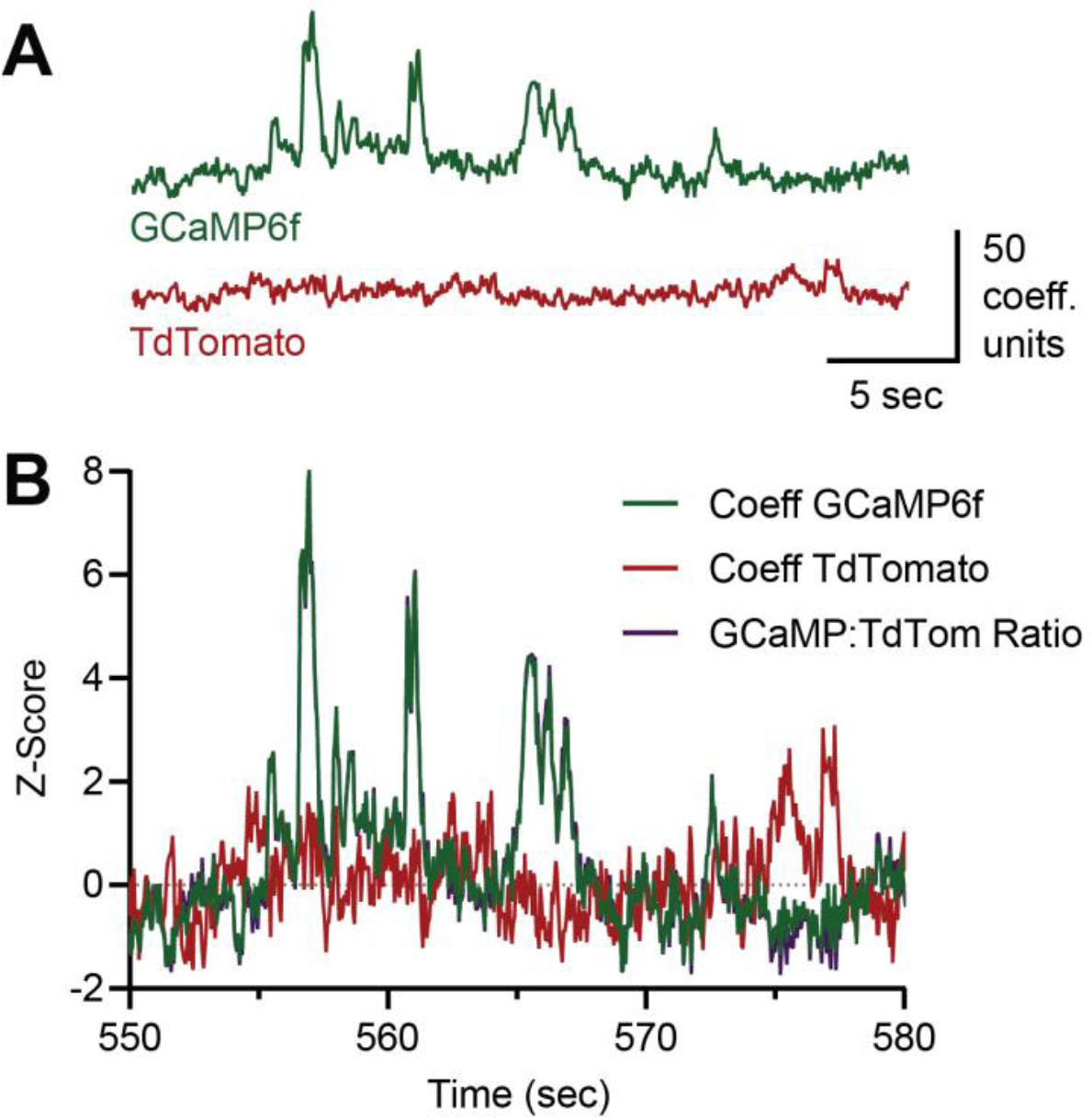
Example fiber photometry recordings of GCaMP6f and TdTomato fluorescence and their ratio. Example spectrally unmixed coefficients of GCaMP6f and TdTomato from the dual-color spectrometer-based fiber photometry system. (B) Z-scores of these example traces, overlayed with their ratio (GCaMP:TdTom), which served as the motion artifact-corrected measure of Ca2+-related changes in fluorescence (as in27) from SST neurons in PL.

**Supplemental Figure 6.**
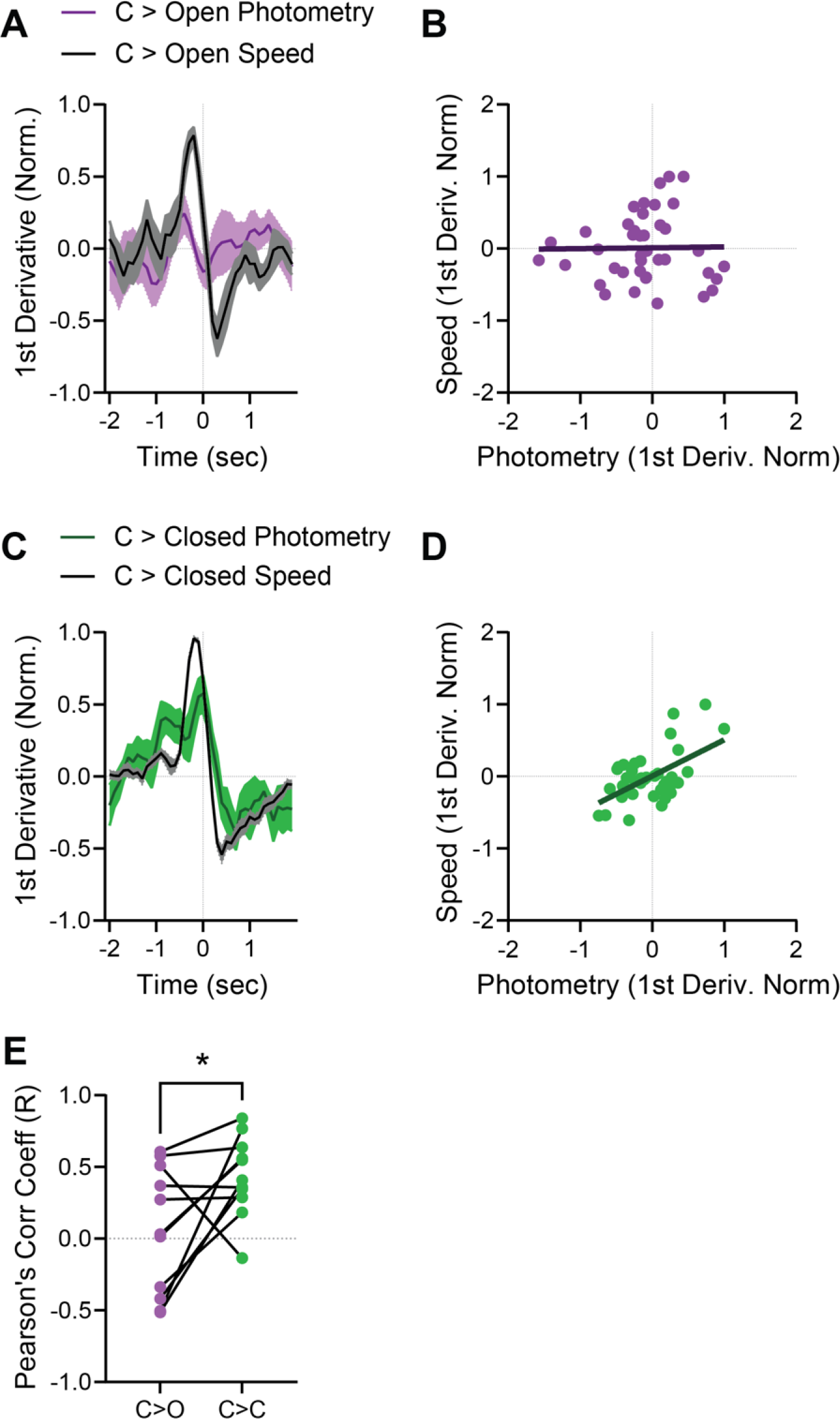
Speed vs. photometry correlations during arm transitions in the EPM. (A) Average normalized 1st derivative of photometry and speed time series across all 11 mice for all C>O transitions. Derivative values normalized to peak value of each mouse in the 4-sec peri-transition time window. (B) Example correlation of 4 seconds of normalized 1st derivative data around C>O transitions for a single mouse. Individual points represent average speed/photometry values at each 0.1-sec interval in the 4-sec transition periods. (C) Average normalized 1st derivative of photometry and speed time series across all 11 mice for all C>C transition. Derivative values normalized to peak value of each mouse in the 4-sec peri-transition time window. (D) Example correlation of 4 seconds of normalized 1st derivative data around C>C transitions for a single mouse. Individual points represent average speed/photometry values at each 0.1-sec interval in the 4-sec transition periods. (E) Pearson’s Correlation Coefficient values for each mouse across both transition types (t10 = 2.40 p < 0.05).

